# Defunctionalizing Intracellular Organelles with Genetically-Encoded Molecular Tools Based on Engineered Phospholipase A/Acyltransferases (PLAATs)

**DOI:** 10.1101/2021.10.10.463806

**Authors:** Satoshi Watanabe, Yuta Nihongaki, Kie Itoh, Shigeki Watanabe, Takanari Inoue

**Affiliations:** Johns Hopkins University School of Medicine, Department of Cell Biology, Baltimore, MD 21205, USA; Johns Hopkins University School of Medicine, Center for Cell Dynamics, Baltimore, MD 21205, USA; Johns Hopkins University School of Medicine, Department of Neuroscience, Baltimore, MD 21205, USA

## Abstract

Organelles vitally achieve multifaceted functions to maintain cellular homeostasis. Genetic and pharmacological approaches to manipulate individual organelles are powerful in probing their physiological roles. However, many of them are either slow in action, limited to certain organelles, or rely on toxic agents. Here, we designed a generalizable molecular tool utilizing phospholipase A/acyltransferases (PLAATs) for rapid induction of organelle defunctionalization via remodeling of the membrane phospholipid composition. In particular, we identified a minimal, fully catalytic PLAAT with no unfavorable side effects. Chemically-induced translocation of the engineered PLAAT to the mitochondria surface resulted in their rapid deformation in a phospholipase activity dependent manner, followed by loss of luminal proteins as well as dissipated membrane potential, thus invalidating the functionality. To demonstrate wide applicability, we then adapted the molecular tool in peroxisomes, and observed leakage of matrix-resident functional proteins. The technique was compatible with optogenetic control, viral delivery and operation in primary neuronal cultures. Due to such versatility, the PLAAT strategy should present a novel utility in organelle biology of diverse contexts.

## Introduction

Membrane phospholipids help cellular organelles acquire distinct properties in their morphology and functionality^1,2^. These membranes not only serve as a physical partition from the cytosol and other organelles, but also become a platform for assembly of functional molecules to play pivotal roles to sustain diverse activities of eukaryotes. There have been experimental approaches to perturb individual organelles. Genetic modification and pharmacological intervention target components in the biogenesis or functional pathways. Despite the utility, their actions are often limited by being either slow in time to allow for compensatory responses in cells, or specific for certain organelles and cannot be readily generalized to other organelles. Other approaches include the use of photosensitizer proteins such as KillerRed^3^, SuperNova^4^ and miniSOG^5,6^ that generate reactive oxygen species (ROS), which oxidize surrounding biocomponents upon light irradiation, leading to defunctionalization of target organelles. Despite their potential in universal application to different organelles, the mode and specificity of action of generated ROS need to be taken into consideration in interpreting experimental results. Another technique utilizes channelrhodopsin which was targeted to organelle membranes to modulate membrane potential^7^. This strategy is exclusively suited for mitochondria that rely on the membrane potential for their functionality.

Due to the fundamental importance of the membranes for organelles to be functional, it is thought that techniques to alter membrane integrity should be effective to investigate the biological significance of individual organelles. With this in mind, we recently engineered actin nucleation promoting factors to generate constrictive force against target organelles for their deformation^8^. Interestingly, when the technique was applied to mitochondria, their major functions such as ATP synthesis were not significantly altered^8,9^. This suggested that induction of deformation is not necessarily sufficient for functional perturbation, and thus radical approaches such as striking remodeling of the membrane composition should be explored.

Phospholipase A (PLA) catalyzes hydrolysis of a phospholipid ester bond at *sn-1* (PLA_1_) or *sn-2* (PLA_2_) position and generates a lysophospholipid and a free fatty acid^10^. Growing evidences have illuminated a close relationship between PLA activity and morphological change of phospholipid membranes. Purified PLA proteins such as PLA_1_, PLA_2_G4A, PLA_2_ from bee venom and transmembrane PLA from bacteria, induced membrane bending, budding, tubulation and pearling in model membranes such as giant vesicles and supported phospholipid bilayers^11–14^. Furthermore, PLA_1_ from snake venom not only swelled isolated mitochondria but also perturbed their functions^15^, while inhibition of PLA_2_ was reported to associate with tubulation of Golgi and trans Golgi network^16^. Of note, these PLAs need a supplementary condition such as non-physiologically high Ca^2+^ concentration for the full catalytic activity. In contrast, one PLA_1_/A_2_ family member, phospholipase A/acyltransferase (PLAAT), has no requirement of such supplementary condition, and was reported to retain unique characteristics to degrade organelle membranes during development of lens transparency^17^.

To develop a molecular tool for organelle defunctionalization that is rapidly inducible, generalizable and target specific, we saw a strong potential in the PLAAT family members. We therefore decided to implement PLAAT proteins in chemically and optically inducible protein dimerization paradigms to reorganize phospholipid composition of individual organellar membranes. After a series of optimizations, we demonstrated that these tools can successfully defunctionalize mitochondria and peroxisomes. The PLAAT molecular tool is genetically-encoded, scalable, and also modularly designed, thus facilitating studies on physiological roles of organelles in various cellular processes.

## Results

### Chemically-inducible translocation of PLAAT family proteins can deform mitochondria through its phospholipase activity

In order to test the concept of utilizing human PLAAT family members for manipulation of morphology and function of organelles, we first chose one of the well-studied members, PLAAT3 (also known as AdPLA, HRASLS3, HREV107, PLA2G16). We then designed an experiment to achieve rapid recruitment of a full-length PLAAT3 to a target organelle using a chemically-inducible dimerization (CID) system (Schema in **Fig. 1a**). Since a previous *in vitro* study showed that PLA deforms isolated mitochondria^15^, we chose mitochondria as a model organelle. To induce translocation of PLAAT3 from the cytosol to the mitochondria surface in living cells, we used the FKBP-FRB dimerizing unit which rapidly and strongly interacts with each other upon rapamycin treatment^18^. We constructed two plasmids (**Fig. 1b**, top) encoding PLAAT3 fused N-terminally to mCherry-FKBP (mCherry-FKBP-PLAAT3) and CFP-FRB fused C-terminally to a mitochondria outer membrane targeting sequence of monoamine oxidase A (CFP-FRB-MoA)^19^. These two constructs were co-transfected into COS-7 cells and subjected to fluorescence live-cell imaging while rapamycin was added to the cells at 100 nM. Within 35 minutes of rapamycin treatment, mitochondria were fragmented and swollen (**Fig. 1b**, upper panels, and **Supplementary Movie 1**). To examine whether this morphological change depends on the phospholipase activity, we repeated the experiment with a lipase-dead (LD) mutant of PLAAT3 where the Cys-113 residue that is critical for the catalytic activity is mutated to serine^20–22^. With mCherry-FKBP-PLAAT3-LD, there was no detectable morphological change in mitochondria (**Fig. 1b**, lower panels, **Supplementary Movie 2**). These results showed that inducible translocation of PLAAT3 to mitochondria membranes can rapidly alter overall morphology through phospholipase activity.

**Figure 1:**
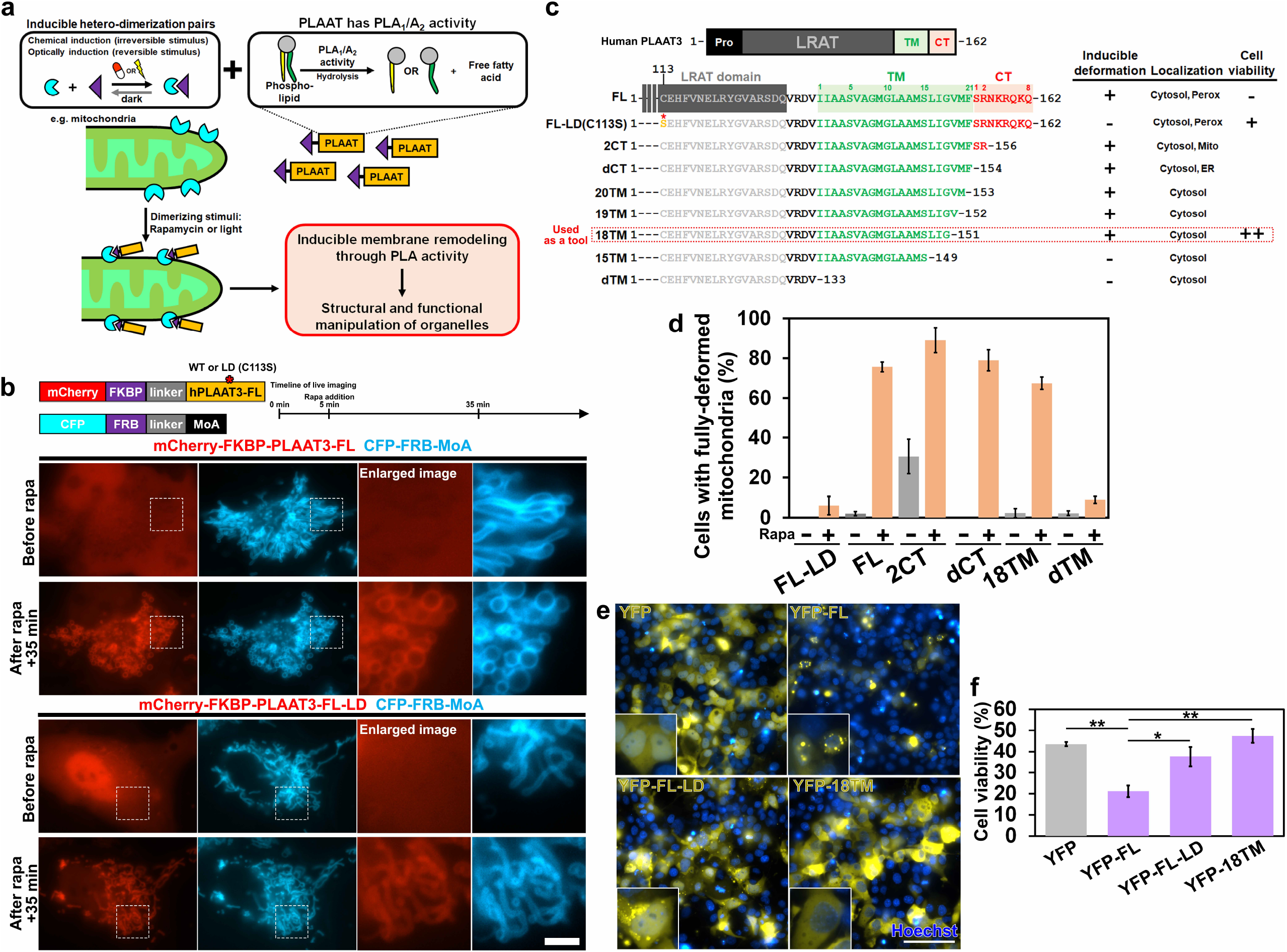
Inducible mitochondrial manipulation by PLAAT3 and its optimization. **a** A conceptual schema describing rapidly inducible manipulation of intracellular organelles based on the integration of chemically- or optically-triggered hetero-dimerization and phospholipid remodeling by PLAAT enzymes. This strategy anticipates rapid switching of the PLAAT activity right at the target membrane-bound organelles (e.g., mitochondria) to induce their deformation as well as defunctionalization. **b** Fluorescence images of COS-7 cells expressing CFP-FRB-MoA (mitochondrial anchor, cyan) along with either mCherry-FKBP-PLAAT3-FL (red) or mCherry-FKBP-PLAAT3-FL-LD (lipase-dead PLAAT3, red) before and 35 mins after addition of 100 nM rapamycin. Enlarged images to highlight mitochondria morphology are shown on the right. Experimental timeline and schematic drawing of constructs used in this experiment are shown on top. **c** A summary of PLAAT3 domain structures (top), as well as C-terminal sequences of full-length PLAAT3 along with its lipase-dead mutant (LD) and a series of truncation mutants (bottom). Properties such as efficiency of inducible deformation, subcellular localization and cell viability were scored based on experiments described below, and indicated for each mutant tested (right). Pro: proline-rich domain, LRAT: Lecithin-retinol acyltransferase (also responsible for PLA_1_/A_2_ activity), TM: putative transmembrane domain, CT: C-terminus domain, FL: full length of human PLAAT3, LD: lipase dead harboring C113S point mutation, 20TM, 19TM, 18TM, and 15TM: PLAAT3 mutant with truncations in TM and a full defect of CT, dTM: PLAAT3 mutant totally defective of TM and CT. **d** A fraction of COS-7 cells indicating fully-deformed mitochondria was calculated before (grey) and after (light brown) rapamycin treatment, and presented for cells expressing mCherry-FKBP-FL-LD, mCherry-FKBP-FL, mCherry-FKBP-2CT, mCherry-FKBP-dCT, mCherry-FKBP-18TM, or mCherry-FKBP-dTM and CFP-FRB-MoA. n = 104, 110, 121, 96, 138, 115, 103, 102, 97, 93, 125, and 111 cells from left to right; analyzed from three individual experiments. **e** Fluorescence images of cells expressing YFP, YFP-FL, YFP-FL-LD, or YFP-18TM, obtained at 48-hr post-transfection. These cells were stained with Hoechst. Insets are zoom-in images. Scale bar = 100 μm. **f** A number of YFP positive cells shown in **e** was calculated and used as a measure of cell viability. n = 591, 556, 499 and 601 cells from left to right; analyzed from three individual experiments. Error bars indicate means ± s.d..

In humans, all 5 PLAAT family members consist of the following domains: Proline-rich (Pro) domain in the N-terminus, lecithin-retinol acyltransferase (LRAT) domain, putative transmembrane (TM) domain and C-terminus region in the most C-terminal end (CT) (**Supplementary Fig. 1a**). The conserved sequences among the PLAAT family motivated us to investigate whether members other than PLAAT3 possess the ability of inducing mitochondria deformation when used in the same translocation scheme described for PLAAT3. As a result, PLAAT1, 2 and 4, but not PLAAT5, induced mitochondrial deformation that is comparable to PLAAT3 (**Supplementary Fig. 1b**). We additionally tested PLAAT4 this time with the truncated TM (PLAAT4-dTM), as this simpler form reportedly retains the phospholipase activity *in vitro*^23^. Unexpectedly, we did not observe mitochondrial deformation (**Supplementary Fig. 1b**). Likewise, the full-length PLAAT5 was incapable of inducing the deformation (**Supplementary Fig. 1b**). Since the TM domain of PLAAT5 is shorter than the other PLAAT TMs by 2-3 amino acids (**Supplementary Fig. 1a**), these results suggested that, besides the catalytic activity, the TM is crucial for the CID-mediated membrane deformation.

### Optimization of PLAAT3 to contain unfavorable characteristics: subcellular localization

PLAAT3 has two characteristics that could unwantedly complicate its use in functional assays. First, PLAAT3 decreases a number of peroxisomes^17,24,25^. Second, constitutive expression of PLAAT3 induces cell death in cancer cells^26,27^. To mitigate the effect on peroxisome biogenesis, we generated several PLAAT3 mutants specifically in the CT domain which binds to a peroxisomal biogenesis factor 19 (PEX19)^25^. We first fused YFP-FKBP to the C-terminus of full-length PLAAT3 (PLAAT3-FL-YFP-FKBP), intending to nullify the interaction between PLAAT3 and PEX19 with steric hindrance. However, PLAAT3-FL-YFP-FKBP failed to induce mitochondria deformation upon recruitment (**Supplementary Fig. 2**). Thus, we next truncated the CT domain (**Fig. 1c**), either partially (2CT) or fully (dCT), and observed their subcellular localization. We specifically introduced truncations to the catalytically inactive LD version (C113S) so that the number of peroxisomes remains intact regardless of their localization. While mCherry-PLAAT3-FL-LD was co-localized with peroxisomes, neither of their truncated counterparts, 2CT or dCT, was detected at peroxisomes (**Supplementary Fig. 3a**). Instead, the 2CT and dCT mutants were unexpectedly mislocalized to mitochondria and endoplasmic reticulum (ER), respectively (**Supplementary Fig. 3b, c**).

Nevertheless, we then quantified the extent of rapamycin-triggered mitochondrial deformation with each of these PLAAT3 mutants, and found that FL, 2CT and dCT, but not FL-LD, worked effectively in inducing robust mitochondria deformation (**Fig. 1d**). As expected, due to the mitochondrial pre-localization, 2CT deformed mitochondria even before rapamycin addition to some extent. Based on dCT mutant, we next designed a series of TM truncation mutants where TM amino acids were truncated (20TM, 19TM, 18TM and 15TM) or in its entirety (dTM) (**Fig. 1c**). When we carried out the CID deformation assay using these mutants, 20TM, 19TM, and 18TM led to mitochondrial deformation comparable to FL, while the remaining mutants such as 15TM and dTM did not (**Fig. 1c, d** and **Supplementary Fig. 4**). The fluorescent images and line scanning analysis showed that these mutants were not localized to organelles such as peroxisomes, mitochondria or ER (**Fig. 1c** and **Supplementary Fig. 3a-c**).

Considering 18TM as a promising candidate for the CID operation, we extensively examined its subcellular localization as a YFP fusion and observed no detectable co-localization with any organelles tested (**Supplementary Fig. 5a-e**). To directly characterize the effect on peroxisome biogenesis, we quantified a number of peroxisomes using mSca-peroxi^28^ as a peroxisomal matrix marker. Accordingly, a number of mSca-peroxi-positive signals decreased in FL-expressing cells, but not in YFP- or YFP-18TM-expressing cells (**Supplementary Fig. 6**). This result suggests that 18TM does not have an effect on peroxisome biogenesis.

### Optimization of PLAAT3 to contain unfavorable characteristics: cell viability

We next aimed to address the remaining unfavorable effect of PLAAT3, namely compromised cell viability which has been described in cancer cell lines^26,27^. Thus, our truncated mutants, such as 18TM that no longer localize to peroxisomes, are predicted to exhibit improved cell viability. We therefore counted the number of cells expressing either YFP, YFP-FL, YFP-FL-LD, or YFP-18TM. At 2 days post transfection, the number of YFP-FL positive cells was significantly smaller (21.1%) than the control YFP positive cells (43.6%) (**Fig. 1e, f**). The observed cell toxicity of YFP-FL was due to the phospholipase activity, as YFP-FL-LD expressing cells survived reasonably well (37.5%). In stark contrast, YFP-18TM retained cell viability (47.4%) to a similar level to the YFP control.

### Characterizations of 18TM as a molecular tool

Membrane damage was reported to recruit PLAAT3 to organelle membranes^17^. We thus examined the extent of stress-induced recruitment of 18TM along with its full-length counterpart and found that 18TM does not exhibit this unfavorable property, likely due to the absence of CT and/or the incomplete TM (**Supplementary Figures 7 and 8**) (see **Supplementary Experiment 1** for experimental details). In addition, we characterized the extent, kinetics and dynamics of mitochondria deformation by CID-induced 18TM recruitment (see **Supplementary Experiment 2** for experimental details), where we observed striking deformation that developed over time with three discernible steps - blebbing, tearing, and shrinkage (**Fig. 2** and **Supplementary Movie 3**). Lastly, since DRP1 protein is indispensable for mitochondria fission^29,30^, we asked if the 18TM-mediated mitochondria deformation such as the observed fragmentation is dependent on DRP1. To address this, we executed the CID operation using 18TM in WT and *Drp1* knockout mouse embryonic fibroblasts^31^ (MEFs). Mitochondrial rounding and fragmentation were similarly observed in both cells (**Supplementary Fig. 9a, b**), indicating that the 18TM action is independent of DRP1. Through the cycles of optimization and characterization, we identified a PLAAT3 truncation mutant, namely 18TM, that is devoid of unfavorable characteristics originally associated with the full-length protein, and generated a molecular tool for synthetic induction of mitochondria deformation that depends on the uncompromised phospholipase activity.

**Figure 2:**
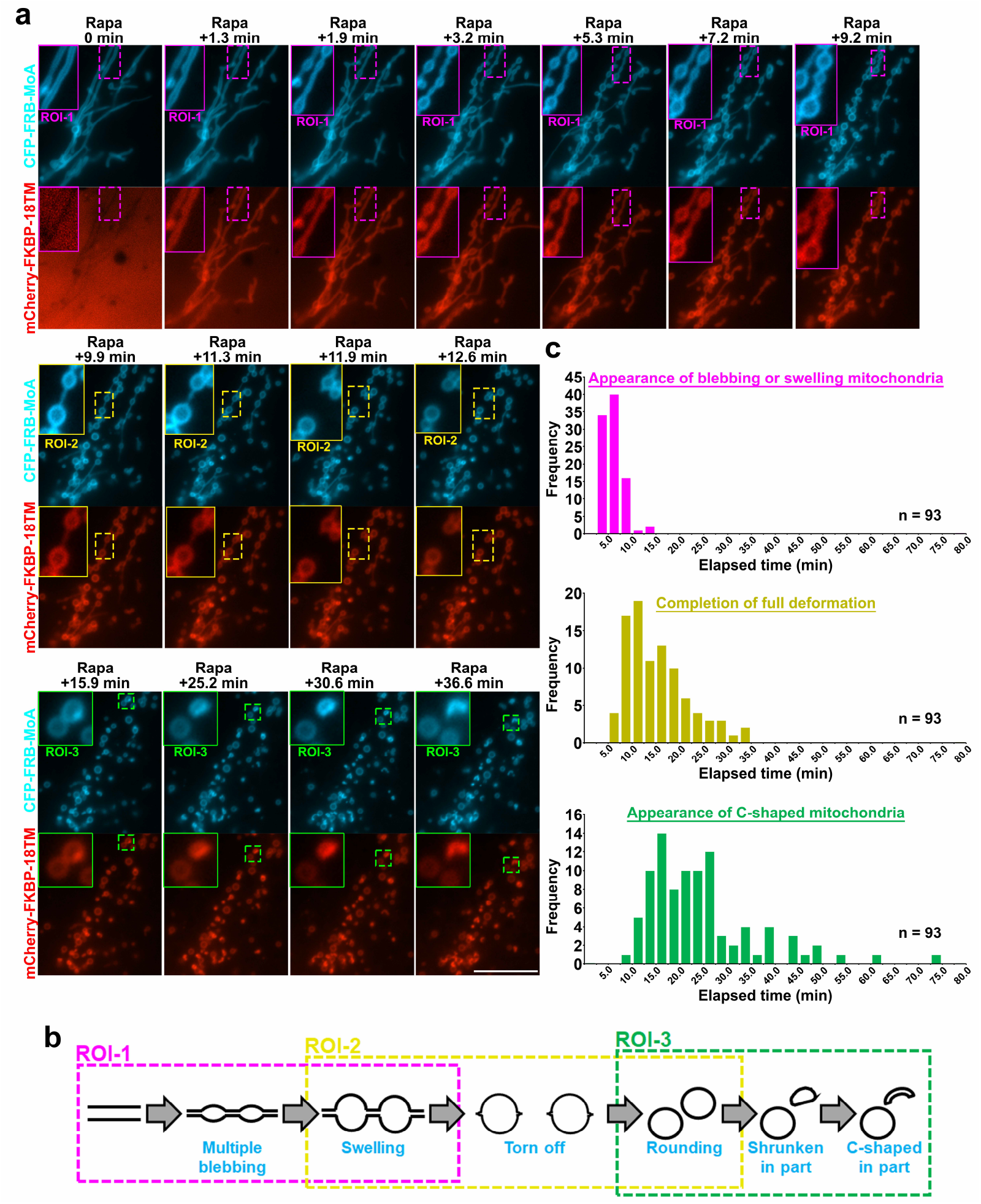
Kinetics of mitochondria morphological changes following induction of 18TM translocation. **a** Fluorescence images of cells expressing CFP-FRB-MoA (cyan) and mCherry-FKBP-18TM (red), indicating three-step morphological changes. ROI-1: Straight-form mitochondria changed into multiple blebbed structures (top panels). ROI-2: Blebs further swelled and finally torn off (middle panels). ROI-3: A part of rounding mitochondria changed into shrunken or C-shaped form (bottom panels). Scale bar = 10 μm. **b** Schematic illustration of the three-step deformation corresponding to the images in **a. c** A number of cells with the indicated deformation type (top: blebbing/swelling, middle: tearing/rounding, bottom: shrinkage/C-shaping) was counted at each time point and presented as a histogram. Ninety-three cells in total were counted from three different experiments. Rapa: rapamycin. ROI: region of interest.

### Defunctionalization of mitochondria following chemically inducible translocation of 18TM

To test whether 18TM-mediated deformation have an influence on mitochondrial functions, we evaluated the two representative properties: membrane potential which empowers ATP synthesis, and autophagic degradation of mitochondria known as mitophagy. We first performed the CID-PLAAT3 experiments to deform mitochondria using YFP-FKBP-18TM (or YFP-FKBP-18TM-LD as a control) together with CFP-FRB-MoA in COS-7 cells. Tetramethyl rhodamine methyl ester (TMRE), an indicator for the mitochondria membrane potential, was then applied at 35 nM for 30 minutes prior to the onset of live-cell imaging. TMRE signals exhibited a moderate decrease 15-30 minutes after rapamycin addition, which continued till they became almost undetectable (**Fig. 3a, b** and **Supplementary Movie 4**). This signal decrease was not due to photobleaching or any systematic artifact, as the TMRE signal intensities were constant in 18TM-LD-transfected cells (**Fig. 3b**). The membrane potential was dissipated even in the presence of 1 mM cyclosporin A, an inhibitor of mitochondrial permeability transition pore (mPTP) (**Fig. 3c**), suggesting that mPTP opening was not the major cause.

**Figure 3:**
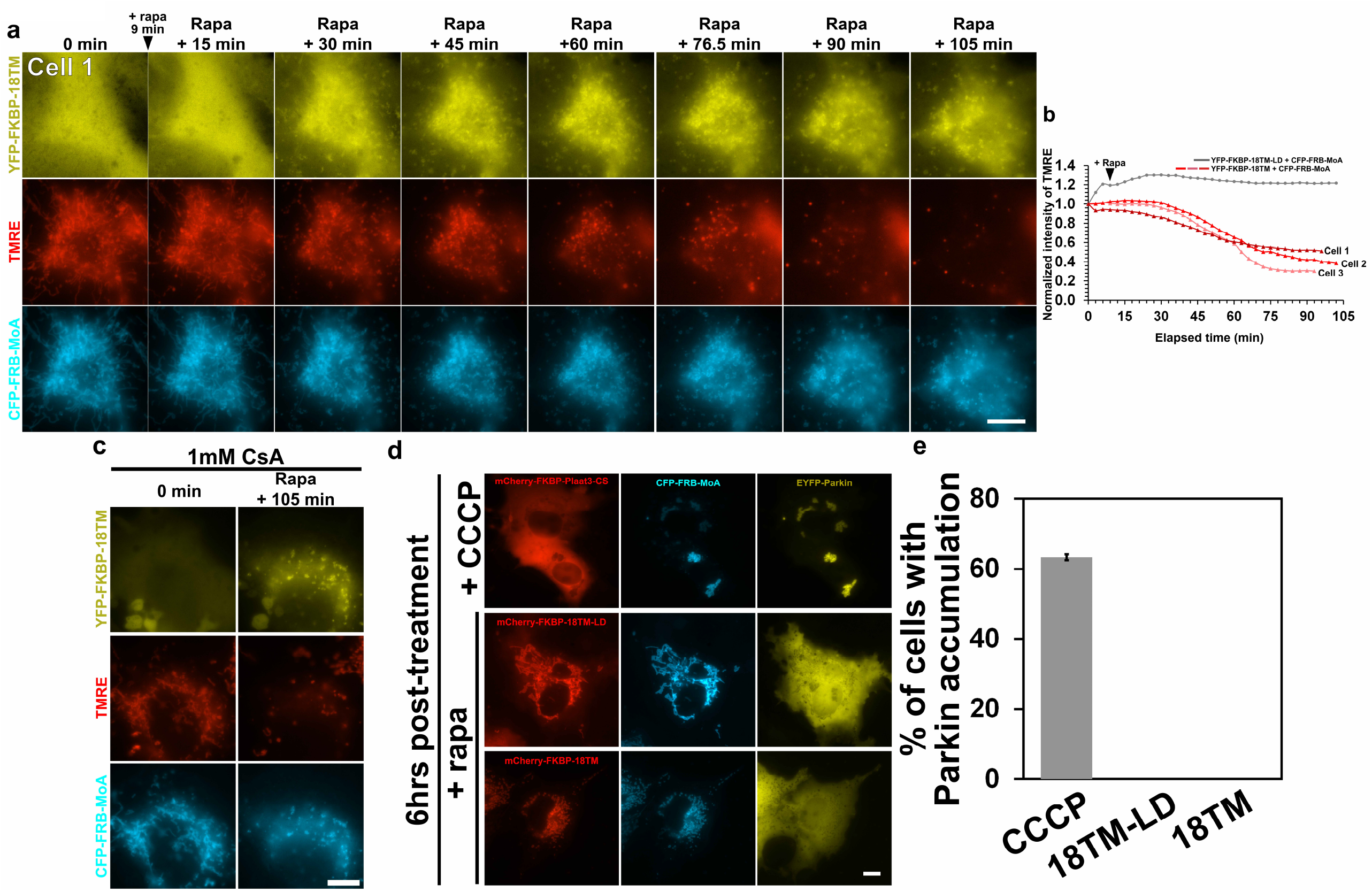
Effect of PLAAT3-18TM-mediated deformation on functional aspects of mitochondria. **a** Time-lapse fluorescence images of TMRE (red) in cells expressing YFP-FKBP-18TM (yellow) and CFP-FRB-MoA (cyan) before and after addition of 100 nM rapamycin. **b** The fluorescence intensity of TMRE in three different cells (Cell 1, 2, 3) was plotted as a function of time after normalized to the value at t = 0. The same measurement was done in cells expressing YFP-FKBP-18TM-LD, instead of YFP-FKBP-18TM, and plotted as a negative control (grey). **c** Fluorescence images of YFP-FKBP-18TM (yellow), TMRE (red) and CFP-FRB-MoA (cyan) in the presence of 1 mM CsA, indicating decreased fluorescence of TMRE 105 mins after addition of 100 nM rapamycin. CsA was added for 1 hr prior to the imaging. **d** Fluorescence images of COS7 cells expressing YFP-Parkin (mitophagy marker, yellow) and CFP-FRB-MoA (cyan) along with either mCherry-FKBP-PLAAT3 (C113S) (red), mCherry-FKBP-18TM (red) or mCherry-FKBP-18TM-LD (red), which were treated with 10 μM CCCP for 6 hrs in cells expressing mCherry-FKBP-PLAAT3 (C113S) (top panels), or with 100 nM rapamycin for 6 hrs in cells expressing mCherry-FKBP-18TM-LD (middle panels) or mCherry-FKBP-18TM (bottom panels). **e** A fraction of cells indicating co-localization of YFP-Parkin and mitochondria was calculated based on the fluorescence images shown in **d**. n = 168, 294 and 241 cells from left to right; analyzed from three individual experiments. Error bars indicate means ± s.d.. Scale bar = 10 μm. Rapa: rapamycin.

We next examined if mitophagy took place following the 18TM-mediated mitochondria deformation. Mitophagy was monitored by accumulation of YFP-Parkin at mitochondria^8^. Cells transfected with mCherry-FKBP-18TM (or mCherry-FKBP-18TM-LD as a control), CFP-FRB-MoA and YFP-Parkin were subjected to 6-hr rapamycin treatment, and the fraction of cells indicating Parkin localization at mitochondria was calculated (**Fig. 3d, e**). Accordingly, we did not observe any sign of mitophagy under 18TM or 18TM-LD conditions, regardless of the striking deformation of mitochondria induced by 18TM. We used carbonyl cyanide 3-chlorophenylhydrazone (CCCP) as a positive control, and as expected, a large fraction of cells (63.4%) accumulated YFP-Parkin at the mitochondria (**Fig. 3d, e**). These data suggest that 18TM triggered loss of membrane potential, but not Parkin-dependent mitophagy.

### Leakage of luminal proteins following mitochondrial recruitment of 18TM

To understand how the remodeling of phospholipids and subsequent deformation led to a loss of the electric potential of mitochondria membranes, we examined the possible leakage of a mitochondria-resident soluble protein, Subunit 9 of mitochondrial ATPase (Su9) in the matrix^32^.

When YFP-FKBP-18TM (or YFP-FKBP-18TM-LD as a control), Su9-CFP and mCherry-FRB-MoA were co-transfected in COS-7 cells, we observed a decrease of Su9-CFP signals upon 18TM-induced deformation (**Fig. 4a**), which appeared to be discrete at each mitochondrion (**Fig. 4b** and **Supplementary Movie 5**). None of these were observed upon recruitment of 18TM-LD (**Fig. 4c** and **Supplementary Movie 6**). A subsequent kinetic analysis of the fluorescence signals at mitochondria further confirmed these results. In particular, while both YFP signals, YFP-FKBP-18TM and YFP-FKBP-18TM-LD, decreased immediately after rapamycin addition and remained constant afterwards, it was only YFP-FKBP-18TM-expressing cells that exhibited a decrease of the Su9-CFP signal upon rapamycin treatment (**Fig. 4d, e**). These results made us suggest that the recruitment of 18TM could increase permeability of the mitochondrial membranes. Accordingly, the previously observed dissipation of membrane potentials (**Fig. 3**) may have been a result of the increased permeability.

**Figure 4:**
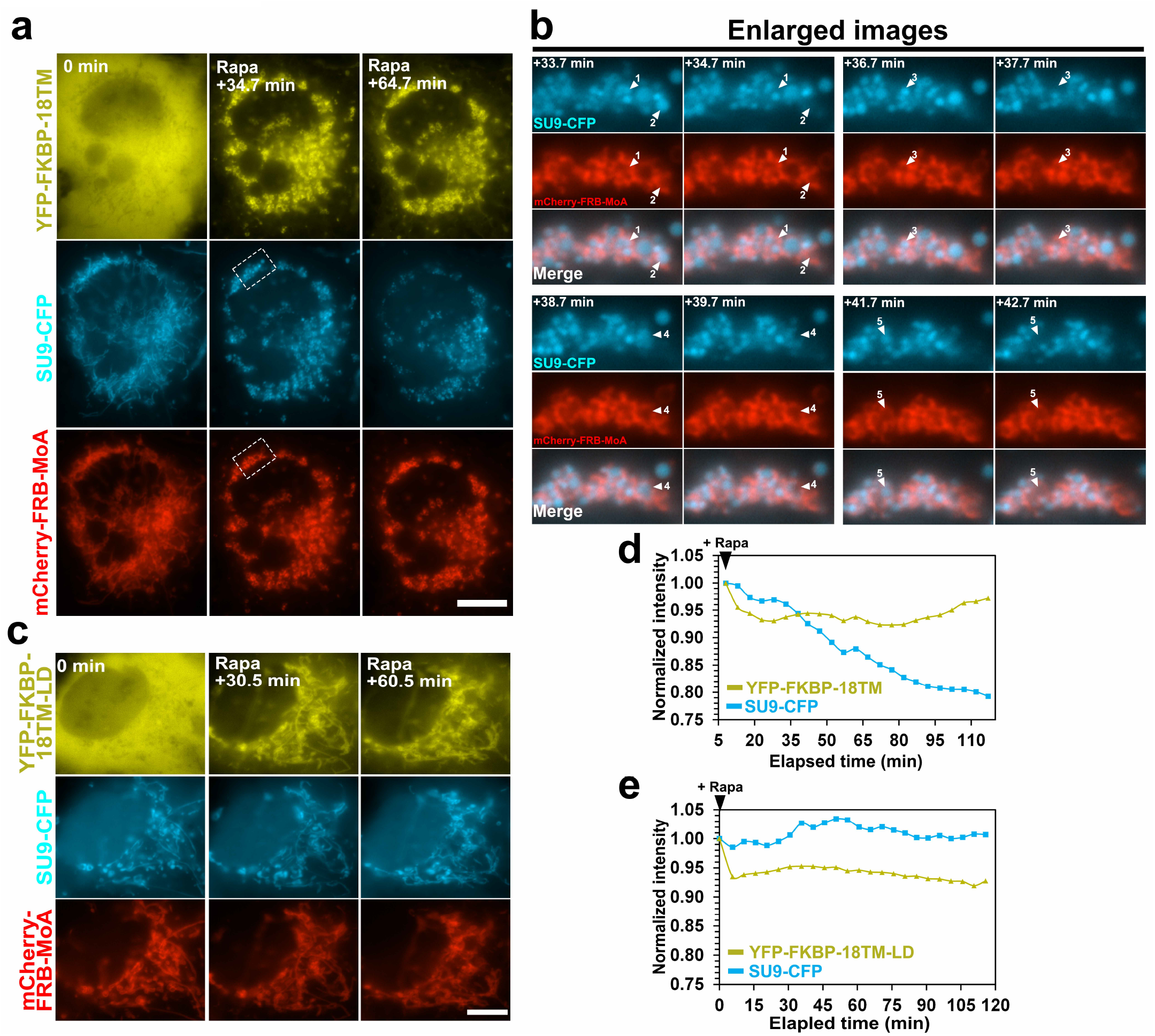
Leakage of luminal proteins following 18TM-mediated mitochondrial deformation. **a-c** Time-lapse fluorescence images of mCherry-FRB-MoA and Su9-CFP (a mitochondria matrix marker), along with either YFP-FKBP-18TM (**a**) or YFP-FKBP-18TM-LD (**b**), in COS-7 cells before and after treatment with100 nM rapamycin. Images indicated by dotted white squares in **a** are magnified in **b. d, e** The fluorescence intensities of Su9-CFP along with either YFP-FKBP-18TM (**d**) or YFP-FKBP-18TM-LD (**e**) were plotted as a function of time after normalized to their own intensity at t = 0. Scale bar = 10 μm. Rapa: rapamycin.

### Optogenetic defunctionalization of mitochondria

To explore the utility of our PLAAT strategy for organelle defunctionalization, we next aimed to achieve an optogenetic operation with 18TM. In particular, we replaced the FKBP-FRB dimerizing pair with an mSspB-iLID pair, which dimerizes upon blue-light exposure. We generated two constructs: 18TM fused to a HaloTag and mSspB (HaloTag-mSspB-18TM) and mitochondria-anchored iLID (iLID-MoA), which were co-transfected in cells along with mCherry-MoA *in trans* to visualize mitochondria. Transfected cells were exposed to intermittent irradiation with blue light (447 nm) for 1 millisecond at a frequency of one minute. Within minutes of the light exposure, we observed that recruitment of HaloTag-mSspB-18TM to the mitochondria (**Fig. 5a**), resulted in deformation similar to what we observed with the CID operation. As a control, we repeated the optogenetic operation with HaloTag-mSspB with no 18TM and found little to no mitochondrial deformation (**Fig. 5b**).

**Figure 5:**
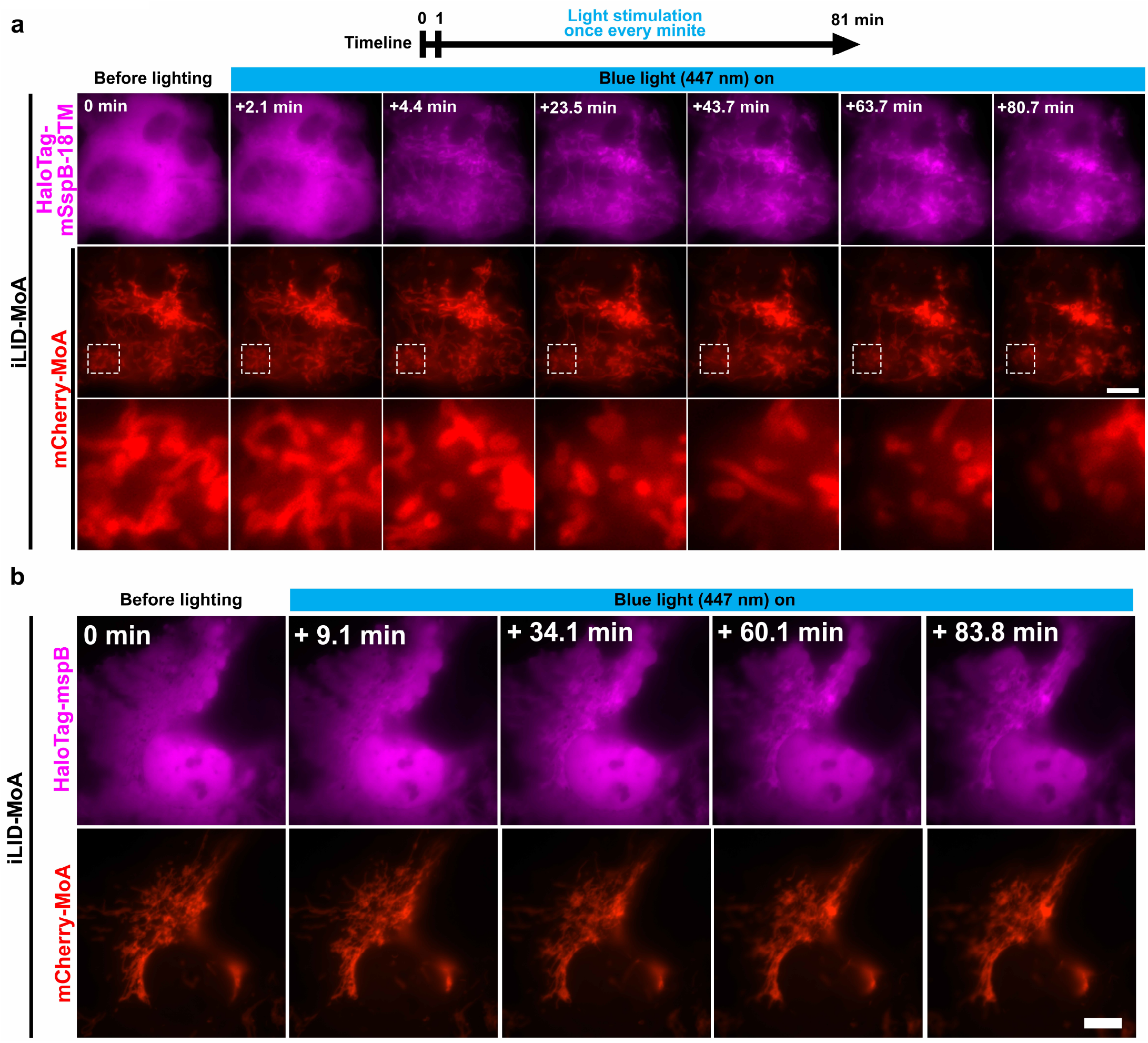
Optogenetic control of 18TM-mediated mitochondrial deformation. **a, b** Time-lapse fluorescence images of COS-7 cells transfected with HaloTag-mSspB-18TM (**a**) or HaloTag-mSspB (**b**) along with iLID-MoA (a mitochondrial anchor) and mCherry-MoA (an indicator of mitochondria morphology). The cells were illuminated by blue light at 447 nm (blue bars on top). The HaloTag-mSspB was visualized with a JF646-conjugated HaloTag ligand which was incubated for 30 mins before an onset of the imaging. Areas marked by dashed-line boxes were magnified at the bottom of **a**. Live-cell imaging was performed in indicated timeline. Scale bar = 10 μm.

### PLAAT-18TM-mediated mitochondria defunctionalization in primary neuronal cultures via viral transduction

We thus far demonstrated the 18TM-mediated mitochondria defunctionalization in cultured cells. Molecular tools like this are useful if applicable to more physiologically relevant samples. We therefore delivered YFP-FKBP-18TM and Tom20-CFP-FRB by adeno-associated virus (AAV) vector into mouse primary hippocampal neurons. More specifically, we raised two types of AAV constructs: YFP-FKBP-18TM (or YFP-FKBP as a control) and Tom20-CFP-FRB, both under the control of a CMV promoter (**Supplementary Fig. 10a**). Tom20 is a component of translocase of the outer membrane (TOM) complex and its N-terminal signal sequence has been interchangeably used to target a protein of interest to mitochondria^19^. Hippocampal neurons were dissected from embryonic mice (E18), then cells were seeded and cultured for 6 days to sufficiently project neurites. At 6 days *in vitro* (DIV6), neurons were infected with two AAVs (YFP-FKBP-18TM and Tom20-CFP-FRB) at multiplicity of infection (MOI) = 40,000 and incubated for 2 days. Subsequent rapamycin treatment triggered deformation of mitochondria, such as dilation, fragmentation and swelling (**Supplementary Fig. 10b**), which was not seen in negative control samples where YFP-FKBP was introduced. These experiments demonstrate applicability of the present PLAAT strategy to different sample preparations beyond culture cell lines.

### Inducible Manipulation of Peroxisomes

To test if the PLAAT strategy can be generalized to other membrane-bound organelles, we chose peroxisomes, since unlike mitochondria there is only a limited number of techniques available for perturbation of peroxisome functions. To direct 18TM to the cytosolic surface of peroxisomes, we made a plasmid encoding PEX3-CFP-FRB, which were transfected in HeLa cells along with YFP-FKBP-18TM (or 18TM-LD as a control) and mSca-Peroxi, followed by live-cell fluorescence imaging. Rapamycin addition successfully recruited PLAAT3-18TM to peroxisomes on a timescale of minutes (**Fig. 6a** and **Supplementary Movie 7**). While we could not detect any noticeable morphological changes under this imaging condition, we observed a decrease in fluorescence intensity of mSca-Peroxi. This decrease was not observed in 18TM-LD-expressing cells (**Fig.6b** and **Supplementary Movie 8**). Tracking individual peroxisomal signals revealed that YFP-18TM and PEX3-CFP-FRB remained constant, while mSca-Peroxi signals were lost sharply and discretely (**Fig. 6c, d**), suggesting that permeability of peroxisomal membranes was increased, similarly to the case of mitochondria. To test if endogenous matrix proteins also leaked out of peroxisomes, we conducted immunocytochemistry in cells expressing YFP-FKBP-18TM (or 18TM-LD) and PEX3-CFP-FRB using an antibody against a catalase, an enzyme which decomposes H_2_O_2_. In 18TM-expressing cells treated with a vehicle (DMSO), catalase was confirmed to localize at peroxisomes (**Fig. 6e**). However, rapamycin treatment in the same cells led to significant decrease in catalase staining (**Fig. 6e, f**). This dissipation of catalase punctate signals did not occur in 18-TM-LD-expressing cells (**Fig. 6e, f**). Catalase-mediated ROS decomposition inside the matrix represents one of the critical functions of peroxisomes. Indeed, mislocalization of peroxisomal catalases to the cytosol often associates with defective *Peroxin* genes that are responsible for peroxisomal biogenesis^33^. Since the 18TM operation led to the loss of such catalases, this particular peroxisomal function must be impaired under this condition.

**Figure 6:**
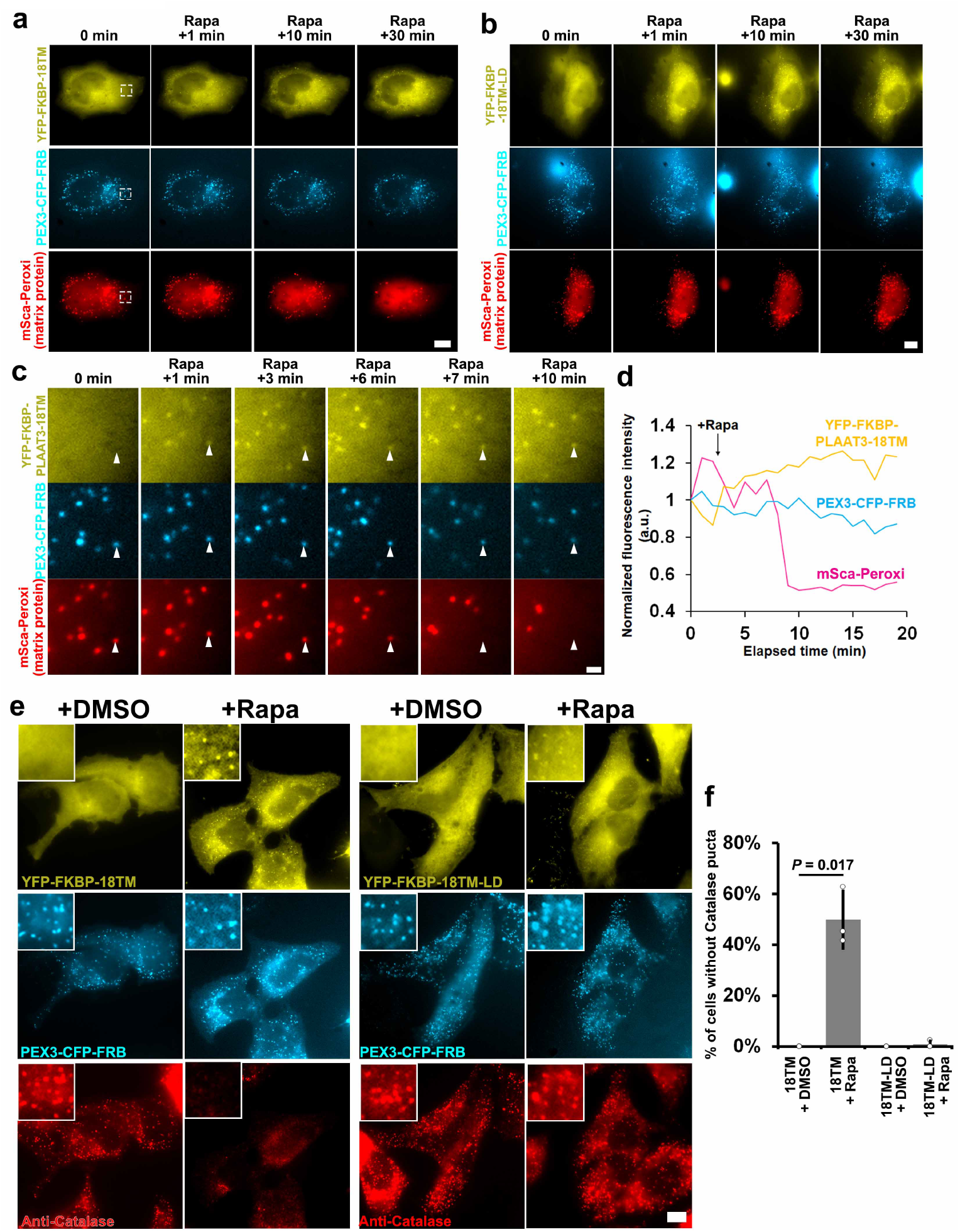
Rapid peroxisome defunctionalization by a chemically inducible PLAAT3-18TM. **a-c** Time-lapse fluorescence images of peroxisomal luminal proteins (mSca-Peroxi) in HeLa cells are presented upon addition of 100 nM rapamycin to trigger chemically-inducible recruitment of PLAAT3-18TM (**a**) or PLAAT3-18TM-LD (**b**). Images indicated by dotted white squares are magnified in **c. d** The fluorescence intensities of YFP-FKBP-18TM, PEX3-CFP-FRB and mSca-Peroxi at a single peroxisome (shown by white arrowheads in c) were plotted as a function of time after normalized to their own values at t = 0. **e** The transfected HeLa cells with indicated constructs were treated with either DMSO or 100 nM rapamycin for 2 hrs before chemical fixation, followed by immunostaining against endogenous catalases. **f** A fraction of cells without catalase signals were calculated based on the fluorescence images in e, where dots and bars indicate individual data points and means ± s.d., respectively (n = 112, 121, 111 and 119 cells from left to right; analyzed from three individual experiments). Rapa: rapamycin. Scale bar = 1 μm for **c** and 10 μm for **a, b** and **e**.

## Discussion

PLAAT members possess phospholipase A/acetyltransferase activity along with promiscuous substrate specificity. With these unique characteristics, PLAAT3 is associated with degradation of almost all types of organelles in eye fiber cells for lens clearing^17^. Despite the strong potential, we realized that PLAAT3 has features that impeded its readily implementation for synthetic operations in living cells. For example, PLAAT3 localizes to peroxisomes and impairs their biogenesis (**Fig. 1, Supplementary Figure 3**) and compromises cell viability (**Fig. 1**). PLAAT3 also gets recruited to mitochondria in response to oxidative and hyperosmotic stress (**Supplementary Figures 7, 8**). Through systematic cell-based characterizations, we generated and assessed a series of PLAAT3 truncation mutants and identified 18TM to retain the desirable membrane deformation ability as much as its full-length counterpart, albeit without the abovementioned undesirable features. Inducible recruitment of 18TM to organelle membranes led to leakage of representative luminal proteins in a manner dependent on the PLA activity. In the case of mitochondria, we also observed the loss of membrane potential, an essential property of mitochondria to galvanize their functions. To demonstrate its utility, we performed the operation with a small chemical and by light, as well as in three different cultured cell lines and primary samples.

One may wonder how the phospholipase A/acyltransferase action by PLAAT on the outer membranes of mitochondria was translated into the leakage of mitochondrial inner proteins such as Su9. There is a protein complex, Ups1-Mdm35, in the mitochondrial intermembrane space that can transfer phospholipids between the two mitochondrial membranes^34,35^. We thus speculated that the lysophospholipids produced at the outer membranes could be routed to the inner membranes, while phospholipids of inner membranes could be simultaneously processed by PLAAT after being transported to the outer membranes. How then does phospholipid remodeling lead to the protein leakage? PLAAT catalyzes a phospholipid molecule to produce a lysophospholipid and a free fatty acid. A loss of one of the two fatty acid chains significantly impacts the membrane stability due to altered physico-mechanical properties^13^, which could be sufficient in increasing membrane permeability to small molecules and proteins likely via membrane ruptures. Since we saw a robust signal of FRB-MoA even after the protein leakage took place (**Figs. 3, 4**), the overall membrane integrity appeared to be retained, and thus these ruptures, if any, may have occurred at a microscopic scale or in a transient manner. Though we dismissed mPTP as a possible pore for the protein leakage (**Fig. 3c**), it is still possible that the remodeled phospholipids could open other transition pores.

In the process of PLAAT3 optimization, we found that a full-length PLAAT3 accumulates at the mitochondria in response to oxidative and hyperosmotic stresses (**Supplementary Figures 7, 8**). Understanding physiological cues that trigger PLAAT3 recruitment to organelle membranes is important^17^. An oxidative insult such as H_2_O_2_ administration leads to peroxidation of membrane phospholipids, which in turn changes the membrane shape and increases the permeability, likely through bursting or pore formation^36,37^. If this applies to mitochondria, the H_2_O_2_ exposure may have caused physical damage to the outer membranes and subsequently induced recruitment of full-length PLAAT3. Of note, hyperosmotic shock can stimulate ROS production in corneal epithelial cells^38^, thus could also induce membrane damages. Hyperosmotic shock may have an alternative or additional role - molecular crowding. Lens fiber cells undergo molecular crowding due to the concentration of crystallin proteins to attain the optimal refractive property^39–42^. Our data implicated that membrane damage and/or molecular crowding could be a physiological trigger of PLAAT3 recruitment to organelles in lens.

An extent of the organelle defunctionalization should be tunable in several different ways. First, the defunctionalization efficiency critically depends on the expression level of each dimerizing unit (FKBP/FRB, and iLID/SspB), so we could adapt promoters of different strength^43^ and other AAV capsid types to achieve an optimal condition. Second, we could adapt and engineer different PLAAT family members or other members of the PLA superfamily from diverse organisms ranging from virus and bacteria to mammals^10^. We expect this alteration to be made feasibly, due to the modular design of our molecular tool (simply replacing 18TM with other PLA_1_ /A_2_ proteins). Besides PLAAT3, PLAAT1, 2, and 4 induced mitochondrial deformation (**Supplementary Fig. 1**). Since PLAAT family members have varying substrate specificity and catalytic strength^10^, it is interesting to apply the same engineering strategy as that for PLAAT3 and develop defunctionalization tools with different characteristics. Since an LRAT protein that catalyzes vitamin A esterification has a very similar sequence and structure to the PLAAT family^44,45^, this enzyme also has a potential to be employed as a molecular tool. Third, the organelle defunctionalization may be achieved by multiplexing molecular tools. For example, our PLAAT tool could be combined with other strategies for organelle manipulation such as those for membrane-bending^8,46^, and those for ROS production^3–5^ where these operations may synergize each other’s action.

Notably, our PLAAT tool is a new addition to a relatively limited pallet of peroxisome research tools. So far, gene modification of *Peroxin* genes and treatment with a catalase inhibitor such as 3-Amino-1,2,4-triazole have been primarily used to investigate physiological roles of this organelle. Since our molecular tool is genetically-encoded, its application should be scalable from single cells to tissues and animals, provided that introduction of a dimerizer molecule or illumination of blue light can be feasibly achieved in these samples. As knockout of *Peroxin* genes in mice associates with deleterious phenotypes including embryonic fatality, it has been challenging to address the physiological role of peroxisomes in mice after development unless conducting a laborious conditional knockout. The present PLAAT-based synthetic operation to induce acute peroxisome defunctionalization in adult mice could thus become as powerful as the conditional knockout, albeit without preparing generations of genetically engineered mice. Once again, the modular design principle should enable expansion of our PLAAT tools beyond mitochondria and peroxisomes. With a collection of these molecular tools, a new type of approaches to organelle biology will become possible.

## Acknowledgements

We would like to thank Hiromi Sesaki for *Drp1* knockout cells and constructs encoding Parkin and Su9, Hideaki Matsubayashi for fruitful discussion, and Hideki Nakamura for technical advice and construct sharing. We also thank the following researchers for constructive discussions on various aspects of the project: Noboru Mizushima, Hideaki Morishita, Tomoya Eguchi, Yukio Fujiki, Kanji Okumoto, Yuuta Imoto, Steve Gould, Michael Wolfgang, Marie Hardwick, Dwight Bergles. We extend our appreciation to Junichi Takagi for kind support of manuscript preparation, and Takeharu Nagai for imaging analysis. This work was supported by discretionary funds to T.I. and the National Institutes of Health (R01GM136858 to T.I.). Sa.W. was supported by a postdoctoral fellowship from the Uehara Memorial Foundation.

## Author contribution

Sa.W. conceived and designed the research, and performed all experiments otherwise mentioned. Sa.W. and T.I. wrote the manuscript. T.I. arranged collaborations. Y.N. performed the peroxisome experiments and contributed to manuscript preparation. Sh.W. and K.I. prepared the primary neuron cultures.

## Competing interests

The authors declare no competing interests.

## Materials and Methods

### Plasmid construction

If not otherwise specified, all plasmids expressing FKBP or FRB were constructed by standard subcloning techniques based on either *pECFP, pEYFP* and *pmCherry* plasmids (Clonetech). Human *Plaat* cDNAs were purchased from ORIGENE [RC208444 for *PLAAT1* (NM_020386), RC212578 for *PLAAT2* (NM_017878), RC200242 for *PLAAT3* (NM_007069), RC201923 for *PLAAT4* (NM_004585), and RC228184 for *PLAAT5* (NM_054108)]. For construction of *pAAV-CMV-YFP-FKBP-18TM, pAAV-CMV-YFP-FKBP-18TM-LD*, and *pAAV-CMV-Tom20-CFP-FRB, pAAV-MCS* (Cell Biolabs) was digested with EcoRI and BamHI, and fragments of *YFP-FKBP-18TM, YFP-FKBP* and *Tom20-CFP-FRB* were inserted to the digested sites using In-Fusion Snap Assembly Master Mix (Takara Bio USA, 638947). To visualize Golgi, lysosomes, autophagosomes, endosomes and nuclear membrane, *Giantin S* and *Lamp1* cDNA were inserted to *pECFP* and cDNAs of *Lc3, Rab5* and *Lamin A* were inserted to *pmCherry* by standard subcloning technique. For PEX3-CFP-FRB, cDNAs encoding human *PEX3, mCerulean3* and *FRB* were inserted into *pcDNA3*.*1(+)* (Invitrogen) vector by standard restriction enzyme subcloning. *mScarlet-SRL*^28^ was obtained from Addgene (#85065). All plasmids were verified by Sanger sequencing.

### Cell culture

HeLa cells, COS-7 cells, HEK293T cells (ATCC) and MEFs were maintained in DMEM (Corning, 10-013-CV) supplemented with 10 % FBS (Sigma Aldrich, F6178), 1 % Pen-Strep (Thermo Fisher, 15140163) at 37 °C under 5 % CO_2_. These cell lines have not been tested routinely for Mycoplasma contamination. In all experiments of chemical dimerization, rapamycin was treated at a final concentration of 100 nM.

### Alignment of amino acid sequences of PLAAT family proteins

Amino acid sequences of human PLAAT family proteins (NP_065119 for PLAAT1, NP_060348 for PLAAT2, NP_009000 for PLAAT3, NP_004576 for PLAAT4, and NP_ 473449 for PLAAT5) were obtained from the NCBI database and aligned by Clustal Omega.

### Cell viability assay

COS-7 cells were transfected with plasmids expressing YFP-FKBP, YFP-FKBP-FL, YFP-FL-LD, YFP-FKBP-18TM. Cells were incubated for 48 hrs and stained with Hoechst33342 (Invitrogen, H3570). Cell pictures were taken using an Eclipse Ti inverted fluorescence microscope (Nikon) equipped with 20× objective lens (Nikon). YFP-positive cells and YFP-negative cells were counted using pictures taken from three different visual fields for each sample and the proportion of YFP-positive cells was calculated as cell viability. Per a visual field, 103-237 cells from three different experiments were counted.

### AAV production and infection

AAV-CMV-YFP-FKBP-18TM, AAV-CMV-YFP-FKBP and AAV-CMV-Tom20-CFP-FRB were produced based on AAV-DJ CMV Expression System (Cell Biolabs, VPK-410-DJ). HEK293T cells were seeded in two six-well cell culture plates (FALCON, 353046) at a density of 1.0□×□10^6^ cells/well and next day transfected with *pAAV-DJ, pHelper* and *pAAV*s encoding transgenes. About 30□mins before transfection, medium change was carried out with 5 % FBS/DMEM supplemented with 1 % Pen-Strep. For one well,□3□μg of a mixture of 1□μg of *pAAV-DJ*, 1□μg of *pHelper*, and 1□μg of *pAAV*s encoding transgenes were used. Six□micrograms of PEI-Max (Polysciences, #24765-1) were used for transfection. After 24□hrs post transfection, the culture medium was changed with 1 % FBS/DMEM supplemented with 1 % Pen-Strep. AAV purification was performed according to previous reports^47,48^. Transfected cells and medium were harvested in one plastic tube 72 hrs after transfection. Three milliliters of chloroform were added to the cell suspension (Sigma, 288306-100ML). To extract AAV particles, the solution was thoroughly vortexed for 5 mins. Cell debris were removed by centrifugation (5 mins, 3,000 × *g*, 4°C). AAV particles were precipitated by adding 9.4 mL of 50 % (v/v) of Polyethylene glycol (PEG) and 7.6 ml of 5M sodium chloride (VWR, SS0430-500GR) and incubating for 2 hrs on ice. AAV precipitates were collected by centrifugation (30 mins, 3,000 × *g*, 4°C). The supernatant was completely discarded and pellets were dissolved with 1.4 ml of PBS. To remove contaminant DNA and RNA, 3.5 μl of 1 M MgCl_2_ (Sigma, M1028), 1.4 μl of DNase I (ThermoFisher, EN0521) and 1.4 μl of 10 μg/μl RNase A (Sigma 10109142001) were added and incubated at 37°C for 20 mins. To remove DNase and RNase, chloroform was applied and solution was thoroughly vortexed for 1 min. By the subsequent centrifugation (5 mins, 12,500 × *g*, room temperature), cell debris were removed and supernatant were collected. This chloroform treatment was repeated at least 3 times. The resultant solution was concentrated in PBS using Amicon ultra centrifugal filter (Millipore, UFC510024). For titration, AAV preparations were treated with 1 unit of DNase I (ThermoFisher, EN0521) at 37 °C for 15 mins, then incubated at 95 °C for 2 mins to inactivate DNase I and finally incubated in 1 % SDS at 70 °C for 15 mins to lyse AAV particles. After optimal dilution of this solution, viral genome copies were quantified by qPCR using SsoAdvanced Universal SYBR Green Supermix (Bio-Rad, 1725272) and primers for a CMV promoter (Forward: 5’-ATAACTTACGGTAAATGGCCCGCC-3’, reverse: 5’-ACTCCACCCATTGACGTCAATGGA-3’)

### Culture and infection of primary mouse hippocampal neurons

Hippocampal neurons from embryonic day 18 (E18) C57/BL6J mice were prepared as previously described^49^. In brief, hippocampi were dissected and incubated in papain for 25 min, then triturated and plated on poly-L-lysine coated 8-well glass chambers (Sigma, P2636, Cellvis, C8-1-N) at a density of 25,000 cells/well in 5 % horse serum (Gibco 26050-088) containing Neurobasal medium (Gibco 21103-049) supplemented with 2 % B-27 (Gibco 17504-044), 2 mM GlutaMax (Gibco 35050-061), and 100 U/ml Penicillin-Streptomycin (Gibco 15140-122). At DIV1, the medium was switched to Neurobasal medium with 2 % B-27 and 2 mM GlutaMax, and neurons were thereafter maintained in this medium. The cultures were stored in a 37 ºC incubator with 5 % CO_2_ until the day of experiments. For the gene transduction into primary neurons, AAVs were infected at MOI = 40,000. Neurons were analyzed 2 days after infection.

### Live-cell imaging

Twenty thousand cells/well for HeLa cells, 40,000 cells/well for COS-7 cells and 30,000 cells/well for MEFs (Wild-type or *Drp*1 KO) were seeded into poly-D-lysine-coated 8 well glass chamber (Sigma, P6407-5MG, Cellvis, C8-1-N). The next day, cells were transfected with mCherry-, YFP-, or HaloTag-FKBP-PLAAT (including PLAAT1, 2, 3, 4, 5, and PLAAT3 mutants) plasmids and FRB-anchor plasmids. The transfection was performed with FuGENE HD or XtremeGENE9 (Sigma, 6365787001) according to the manufacturer’s protocol. Then cells were incubated for 15-24 hrs before live cell imaging. For live cell fluorescence imaging, an Eclipse Ti inverted fluorescence microscope equipped with 60× (for peroxisome observation, Nikon) or 100× (the other observation, Nikon) oil-immersion objective lens and pco.edge sCMOS camera (for peroxisome observation, PCO) or Zyla 4.2 plus CMOS camera (for the other observation, Andor) was used. Time-lapse imaging was conducted at 37 °C, 5 % CO_2_ and humidity using a stage top incubator (Tokai Hit) and done at 0.5-1 min intervals for 60-90 min. For the induction of FKBP-FRB interaction, cells were treated at a final concentration of 100 nM rapamycin. For optical control of inducible 18TM recruitment, COS-7 cells were transfected with plasmids expressing HaloTag-mSspB-18TM (or HaloTag-mSspB), iLID-MoA and mCherry-MoA. Next day, to visualize Halo-Tag-conjugated proteins, JF646-conjugated Halo Tag ligand (Promega, GA1120) was added to culture at a final concentration of 200 nM and cells were incubated for 30 mins before live-imaging. Images were taken using an Eclipse Ti with similar settings. After the first frame of images was taken, blue light (447 nm) started to be irradiated at approximately 1 min interval for 1 millisecond at the lowest stimulus intensity. Imaging was performed for about 80 mins.

### Observation of peroxisome reduction following PLAAT3 expression

To observe changes of peroxisomal numbers, COS-7 cells were transfected with plasmids expressing YFP, YFP-FL or YFP-18TM and mSca-Peroxi. After 48 hrs, imaging was carried out using Eclipse Ti.

### Quantification of mitochondrial deformation and PLAAT3 translocation

For mitochondrial deformation, COS-7 cells were transfected with plasmids encoding mCherry-FKBP-PLAAT3 or -PLAAT3-mutants and CFP-FRB-MoA. Next day, transfected cells were treated with rapamycin for 50 mins. Cell images were taken by 100× oil-immersion objective lens and Eclipse Ti, 93-138 cells from three different experiments were analyzed. Proportion of cells with fully-deformed mitochondria was calculated.

To quantify translocation of FL and 18TM after treatment with agents inducing membrane damage, oxidative stress and hyperosmotic stress, we carried out transfection into COS-7 cells with plasmids encoding mCherry-FKBP, mCherry-FKBP-FL or mCherry-FKBP-18TM (or their YFP versions and LD versions) and CFP-FRB-MoA. Next day, 1 mM of LLOME (Cayman, 16008) was applied to induce membrane damage. For inducing oxidative stress, 100 μM and 500 μM of H_2_O_2_ (Sigma, H1009) were added to culture. For inducing hyperosmotic stress, 200 mM and 500 mM of Sucrose (Sigma, S0389) or 50 mM and 100 mM of NaCl (Sigma, S3014) were applied to culture. These reagents were diluted in 10 % FBS/DMEM supplemented with 1 % Pen-Strep. One (for LLOME treatment) or two (for treatment with the other agents) hrs after addition, cell images were taken using 60× oil-immersion objective lens and Eclipse Ti and 305-405 cells from three different experiments were analyzed. Proportion of cells with merged signals between YFP or mCherry and CFP-FRB-MoA was calculated.

### Imaging analysis

For co-localization analysis, COS-7 cells were transfected with plasmids encoding YFP-, YFP-FL, YFP-18TM (or their mCherry versions) and organelle marker proteins: PEX3-YFP for Peroxisomes, CFP-MoA for mitochondria, CFP-SEC61B for ER, Giantin-CFP for Golgi, LAMP1-CFP for lysosomes, mCherry-LC3 for autophagosomes, mCherry-Rab5 for endosomes, and mCherry-Lamin A for nucleus. In particular, to clearly see autophagosomes, cells were treated with 100 μM of chloroquine for one overnight. Next day, cells were imaged. To quantitatively analyze co-localization, line scanning analysis was performed using NIS element (Nikon). For the intensity analysis of fluorescent proteins and TMRE, regions of interest (ROIs) were manually drawn on analyzed organelles guided by anchor protein fluorescence. Using ROIs, the fluorescence intensities of each color channel were measured using NIS element. For membrane potential measurement using TMRE, COS-7 cells were transfected with plasmids encoding YFP-FKBP-18TM (or YFP-FKBP-18TM-LD) and CFP-FRB-MoA, and next day, were treated with 35 nM TMRE (final concentration) for 20 minutes before live imaging. Imaging was carried out for 90 mins. As a control, FCCP [Carbonyl cyanide 4-(trifluoromethoxy) phenylhydrazone] was added to culture at a final concentration of 10 μM and we confirmed that TMRE signals were quickly lost within 5 mins after the treatment. To examine dependency of loss of membrane potential on mPTP opening, cells were treated with 1 mM CsA for 1hr before imaging. After rapamycin treatment for 105 mins, cells were imaged by Eclipse Ti. While imaging, CsA was not removed. COS-7 cells were also transfected with plasmids encoding YFP-FKBP-18TM, mCherry-MoA and Su9-CFP. Next day, live cell imaging was performed. In quantification of these experiments, the intensities were normalized by the intensity from the first frame. In these experiments, to reproducibly observe protein leakage, we selected and imaged cells with higher expression levels of both 18TM and anchor proteins. To quantify the percentage of catalase-negative cells, cells expressing both YFP-FKBP-PLAAT and PEX3-CFP-FRB were selected. Because overexpression of PEX3-CFP-FRB at the highest level induced its mislocalization to mitochondria, the cells showing mislocalized PEX3-CFP-FRB were excluded from the analysis. The analyzed cells without colocalization between PEX3 and endogenous catalase were counted as catalase-negative cells. The intensities were normalized as the intensity at t = 0 to 1. Cells (111-121 cells) were chosen from three different experiments.

### Immunofluorescence

HeLa cells were seeded at a density of 20,000 cells/well into poly-D-lysine-coated 8 well glass chamber. The next day, cells were transfected with plasmids encoding YFP-FKBP-PLAAT and PEX3-mCeru3-FRB. The transfection was performed with FuGENE HD according to the manufacturer’s protocol. After 24 hrs post transfection cells were treated with 100 nM rapamycin or DMSO for 2 hr. Then cells were washed with PBS (not containing Mg^2+^ and Ca^2+^) thrice and fixed with 4 % paraformaldehyde (PFA, Electron Microscopy Sciences, 15714) in PBS for 10 min at room temperature. Next, cells were washed with PBS thrice and permeabilized and blocked with 2 % bovine serum albumin and 0.1 % Triton X-100 in PBS for 1 hr at room temperature. After blocking, cells were incubated with primary antibody against Catalase (Cell Signaling Technology, D4P7B, 1:1,000) diluted in blocking buffer overnight at 4 °C. Cells were then washed with PBS thrice and incubated with diluted anti-rabbit secondary antibody conjugated with Alexa Fluor 594 (Thermo Fisher Scientific, A-11072) at 1:1,000 for 1 h at room temperature. Cells were washed with PBS thrice again and subjected to fluorescence imaging. The fluorescence imaging was performed on an Eclipse Ti inverted fluorescence microscope equipped with 60× oil-immersion objective lens and pco.edge sCMOS camera.

### Statistical information

Statistical parameters including the definition and exact values of n (number of cells and experiments) and deviation are described in figures and corresponding legends. Data are represented as mean ± s.d.. *t*-tests were performed using Microsoft Excel.

## Notes

### Competing Interest Statement

The authors have declared no competing interest.

