## Supplementary Figures 1-10 for "Defunctionalizing Intracellular Organelles with Genetically-Encoded Molecular Tools Based on Engineered Phospholipase A/Acyltransferases (PLAATs)"

#### Supplementary Figure 1

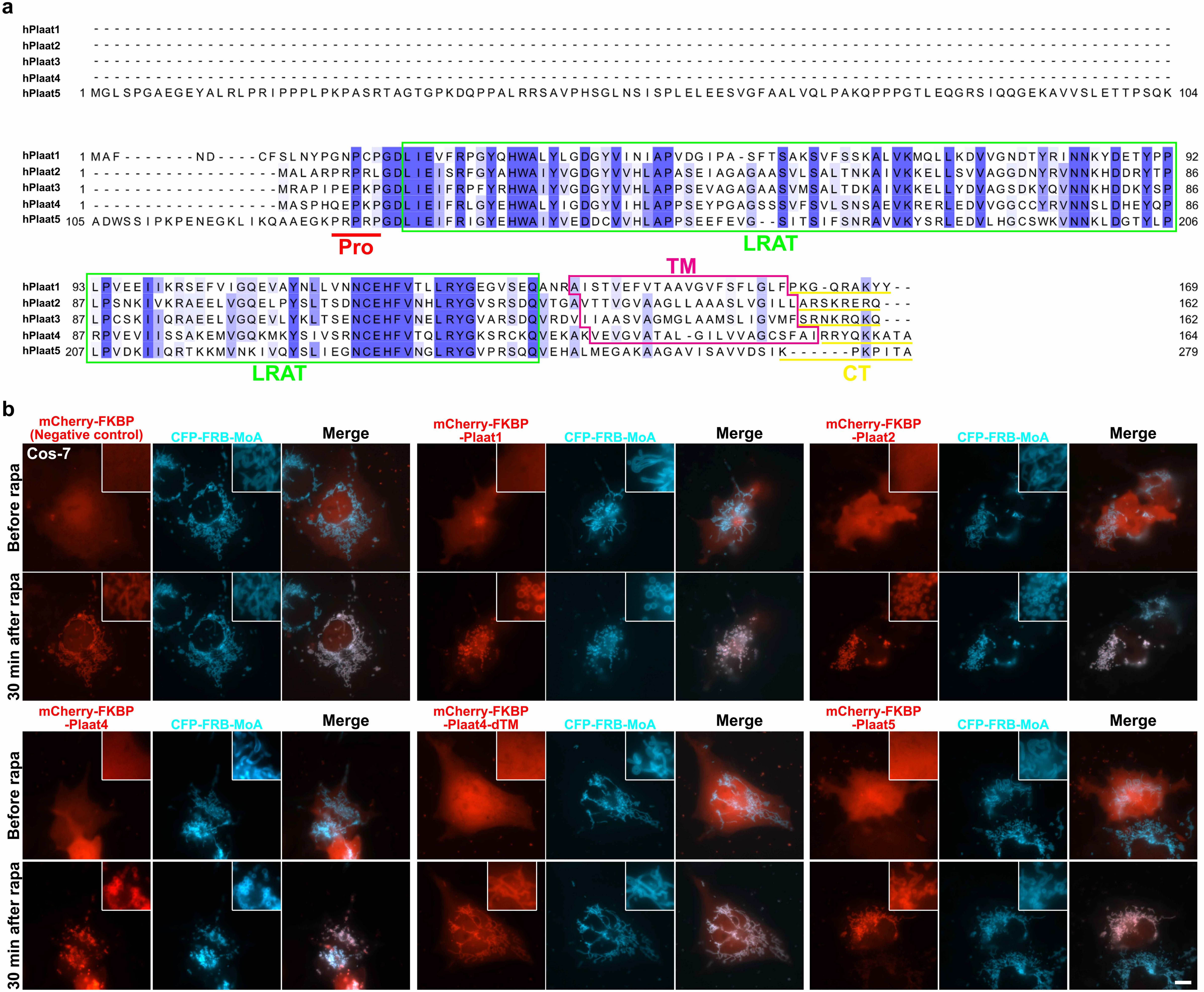

### Supplementary Figure 2

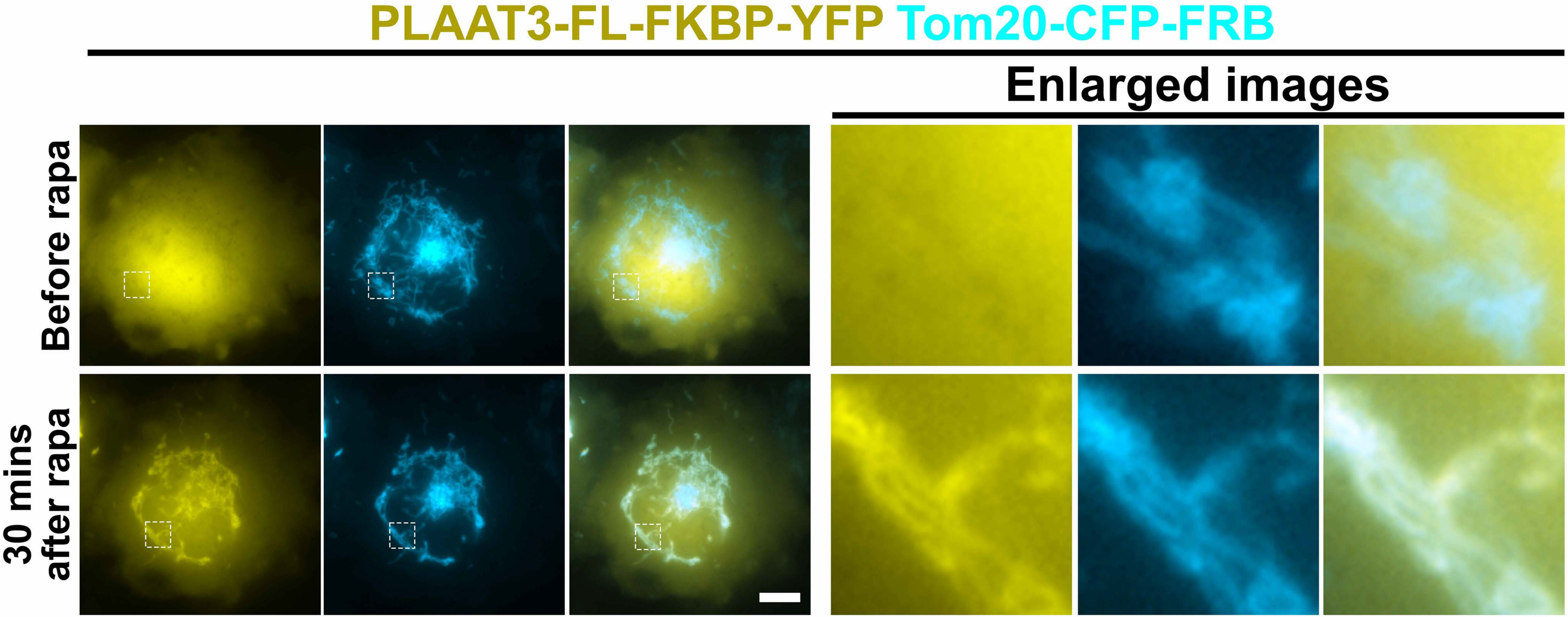

### Supplementary Figure 3

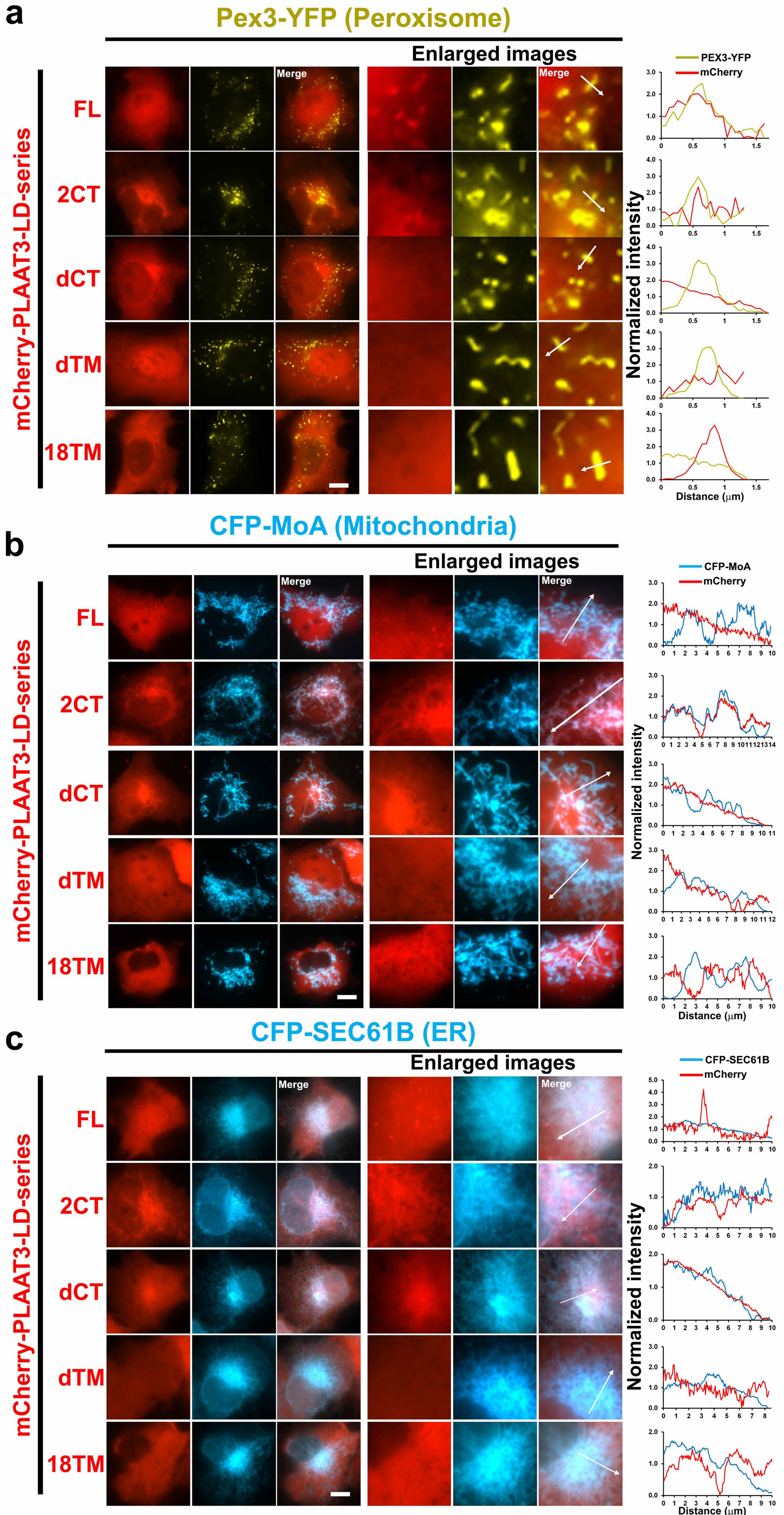

### Supplementary Figure 4

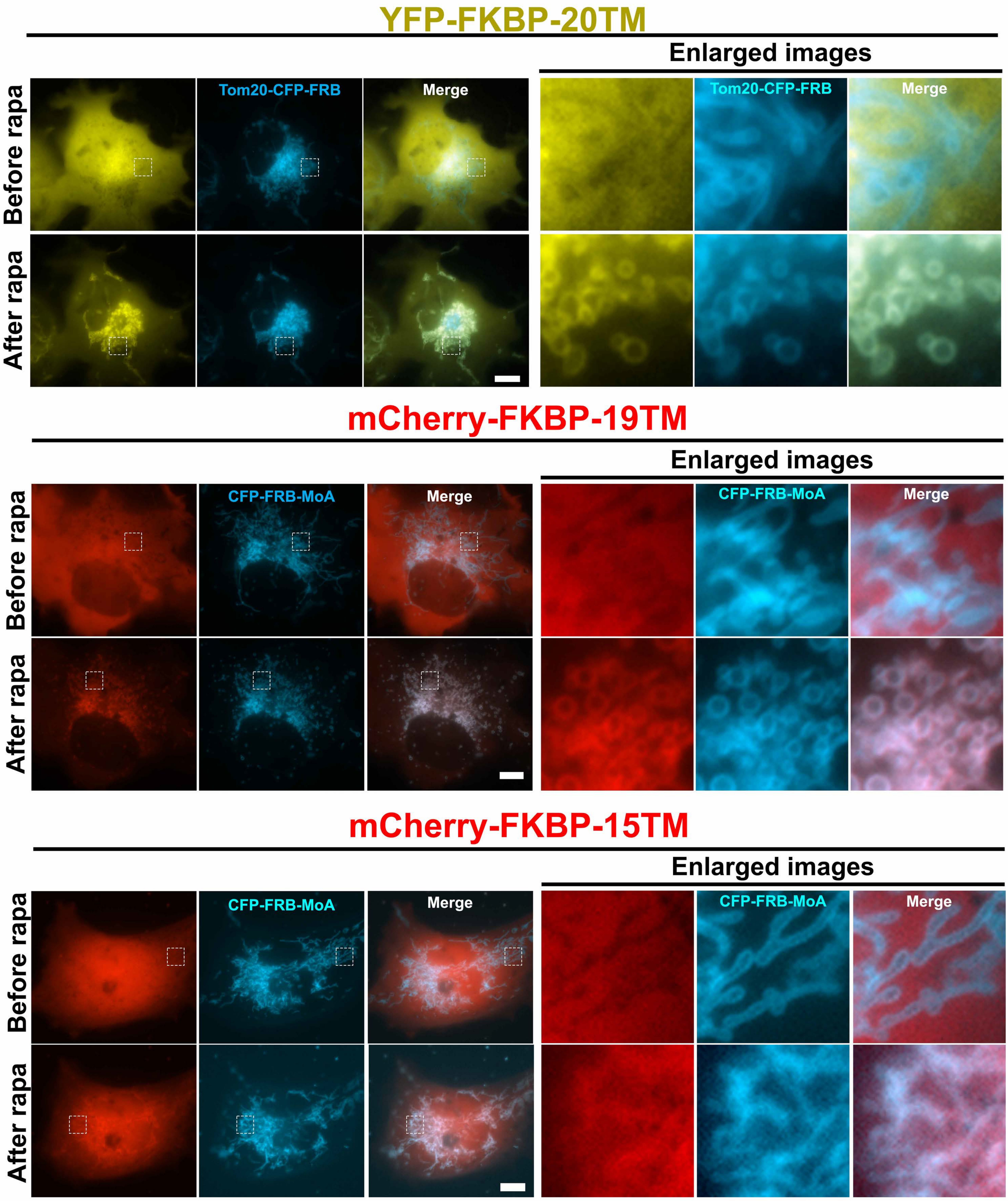

### Supplementary Figure 5

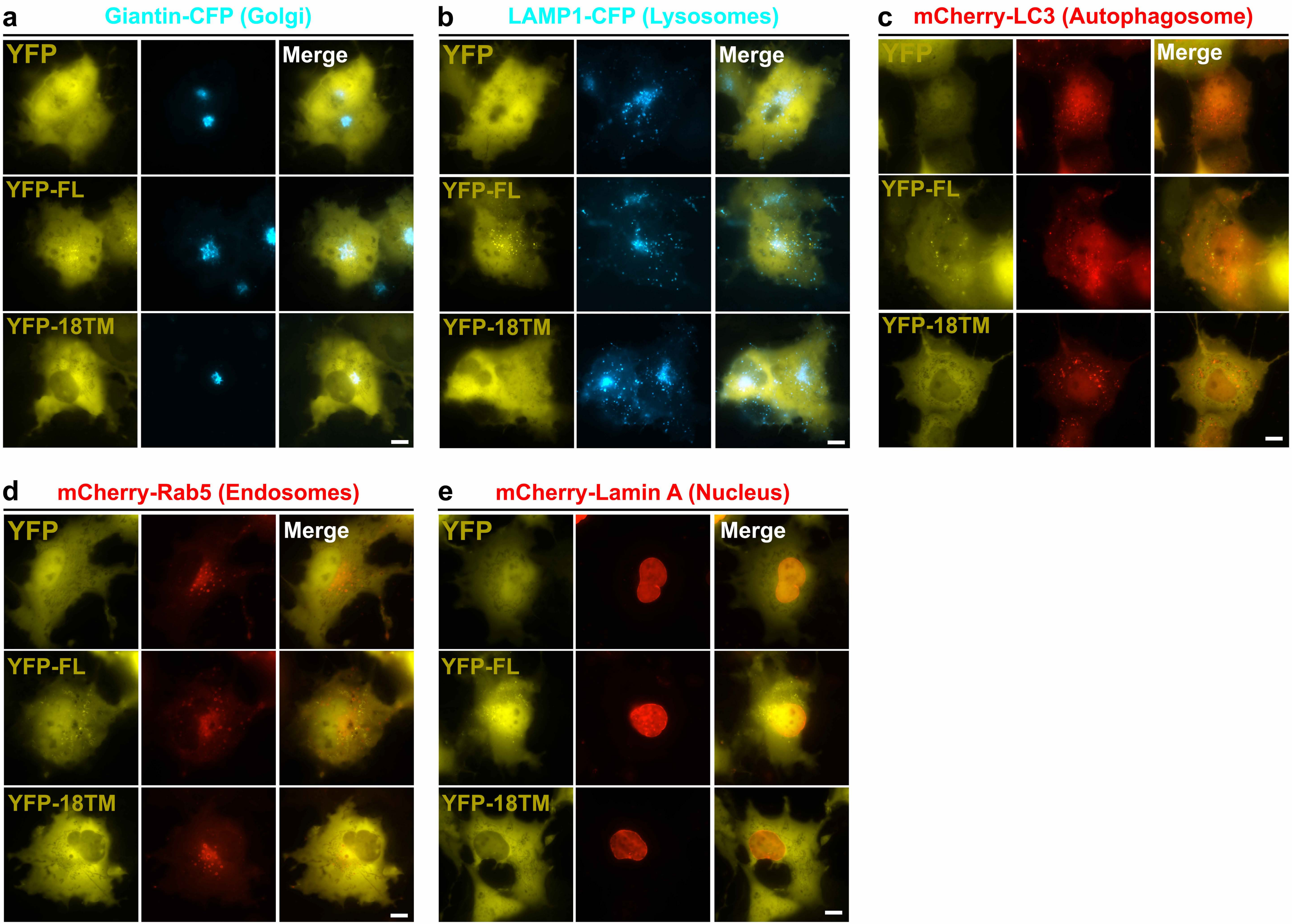

### Supplementary Figure 6

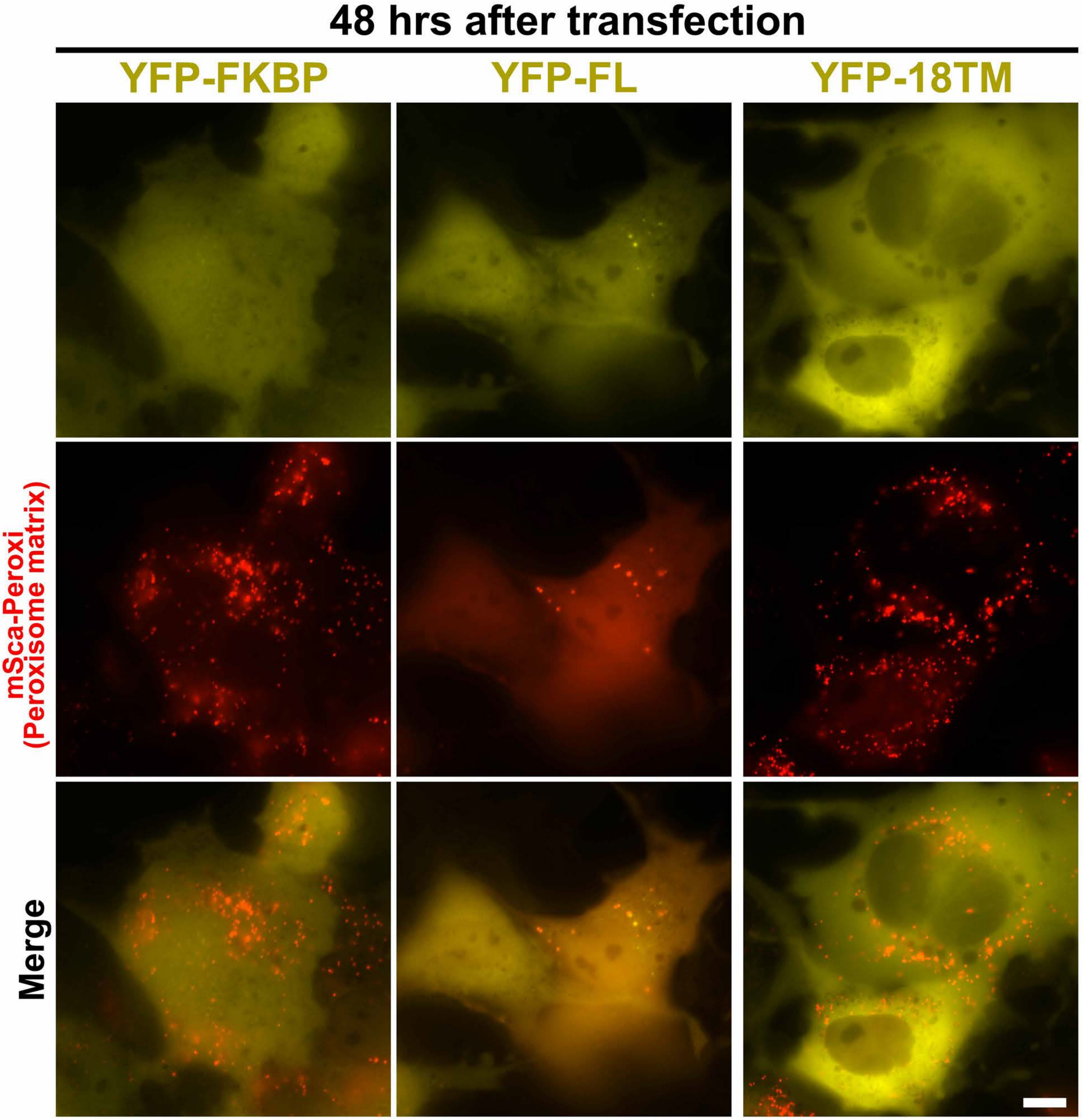

### Supplementary Figure 7

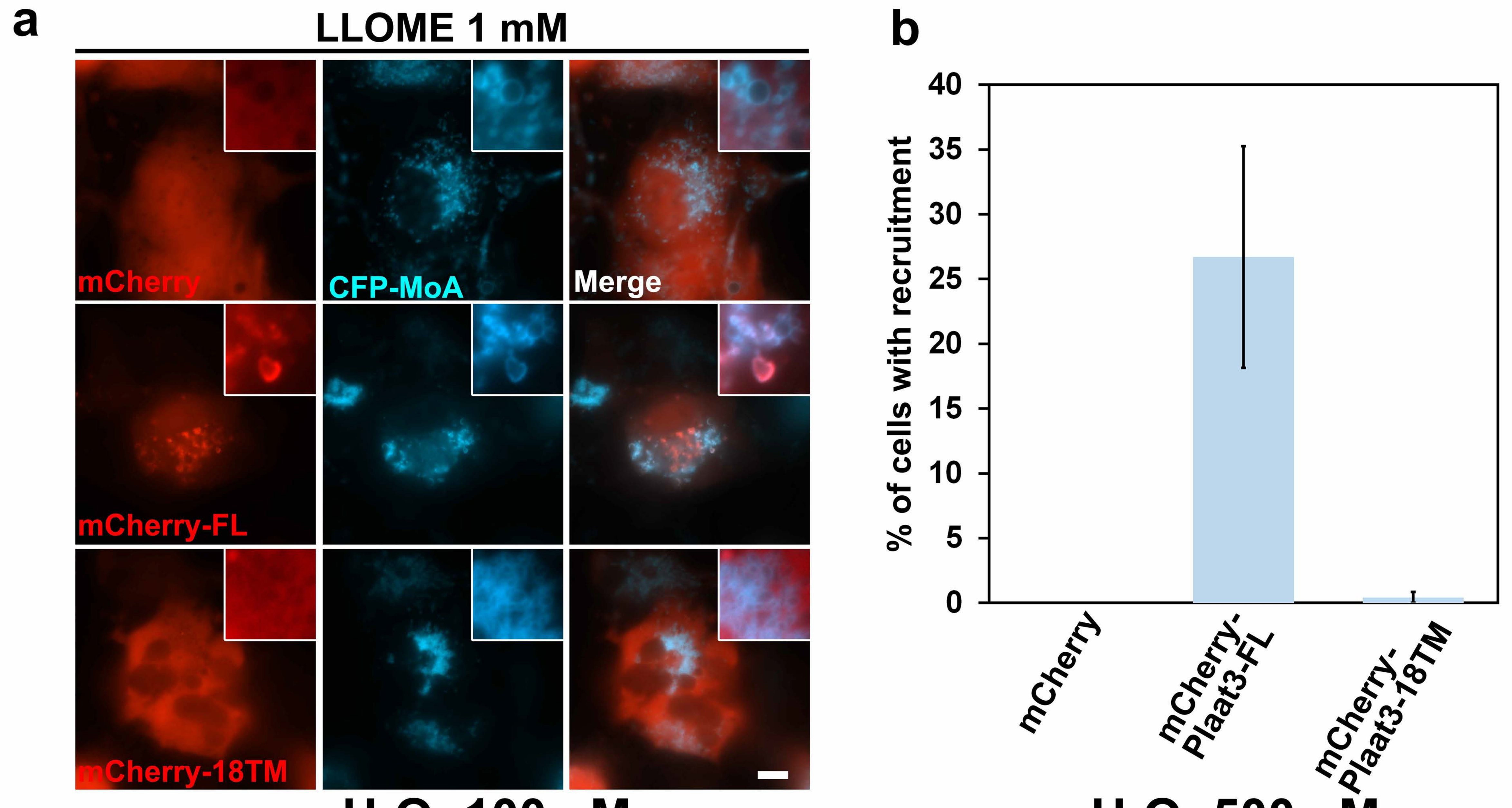


i

| | Spontaneous recruit | Stress-induced recruit |
|---|---|---|
| FL | + | + |
| FL-LD | + | + |
| 18TM | - | - |
| 18TM-LD | - | - |

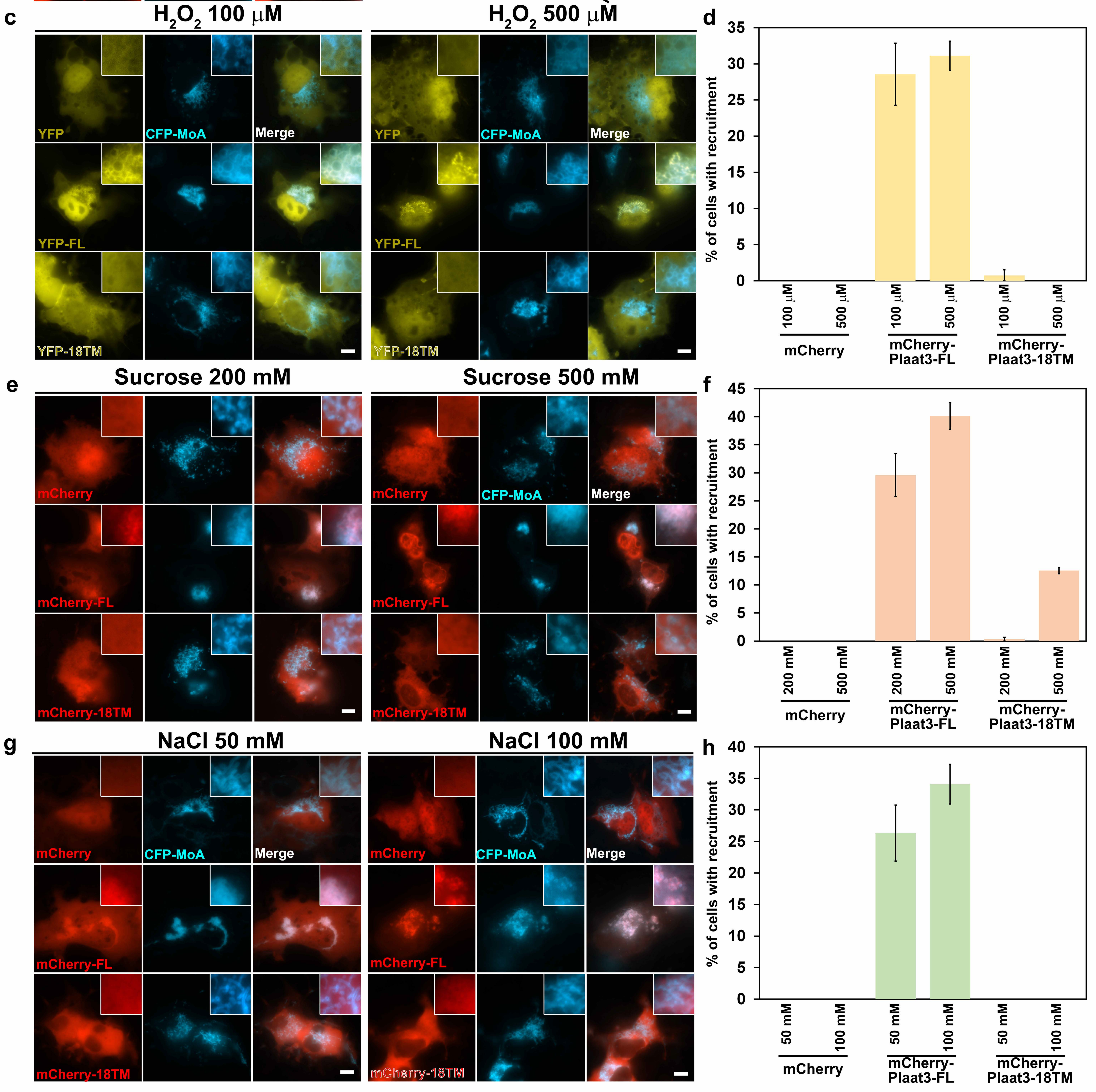

### Supplementary Figure 8

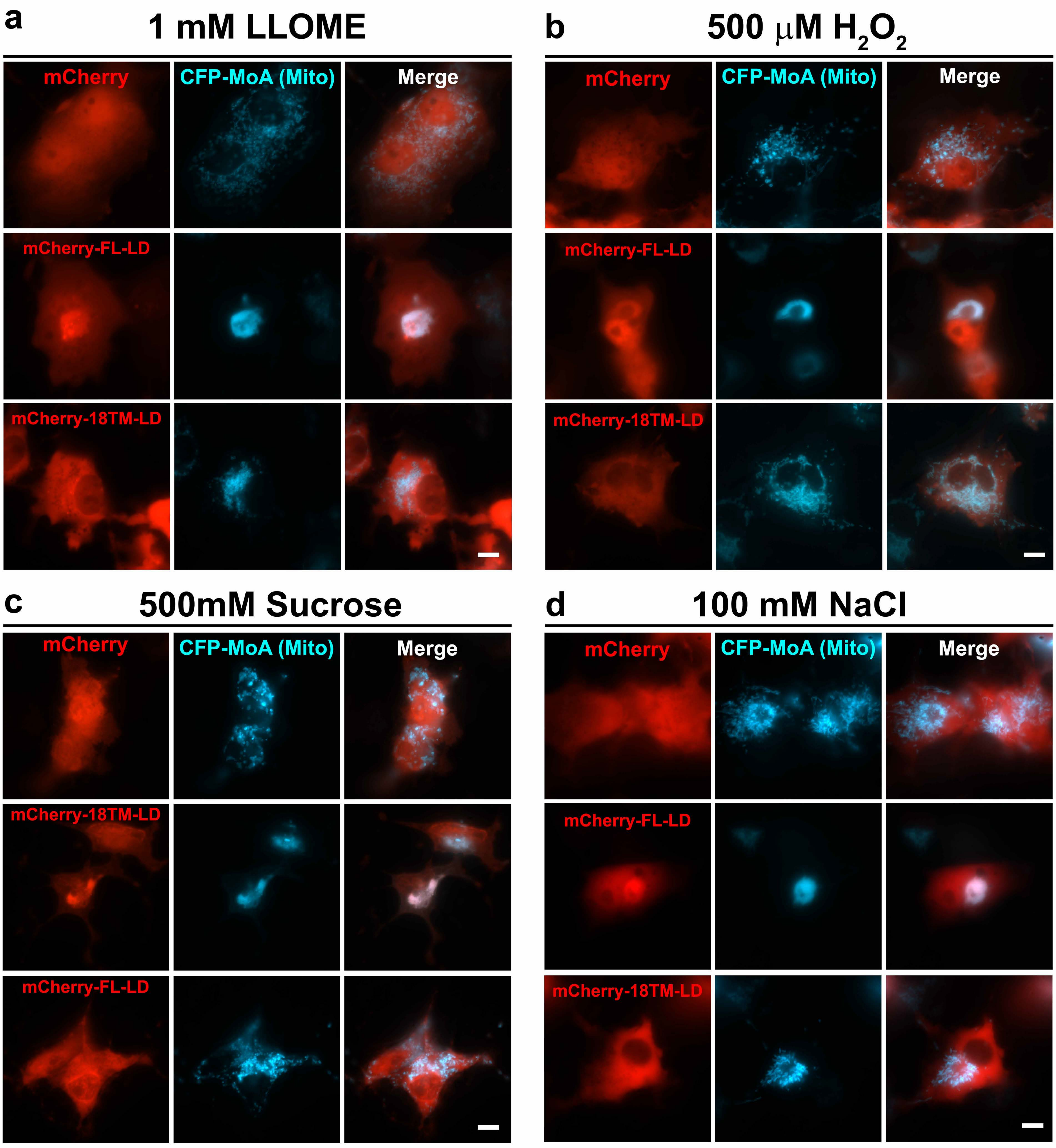

### Supplementary Figure 9

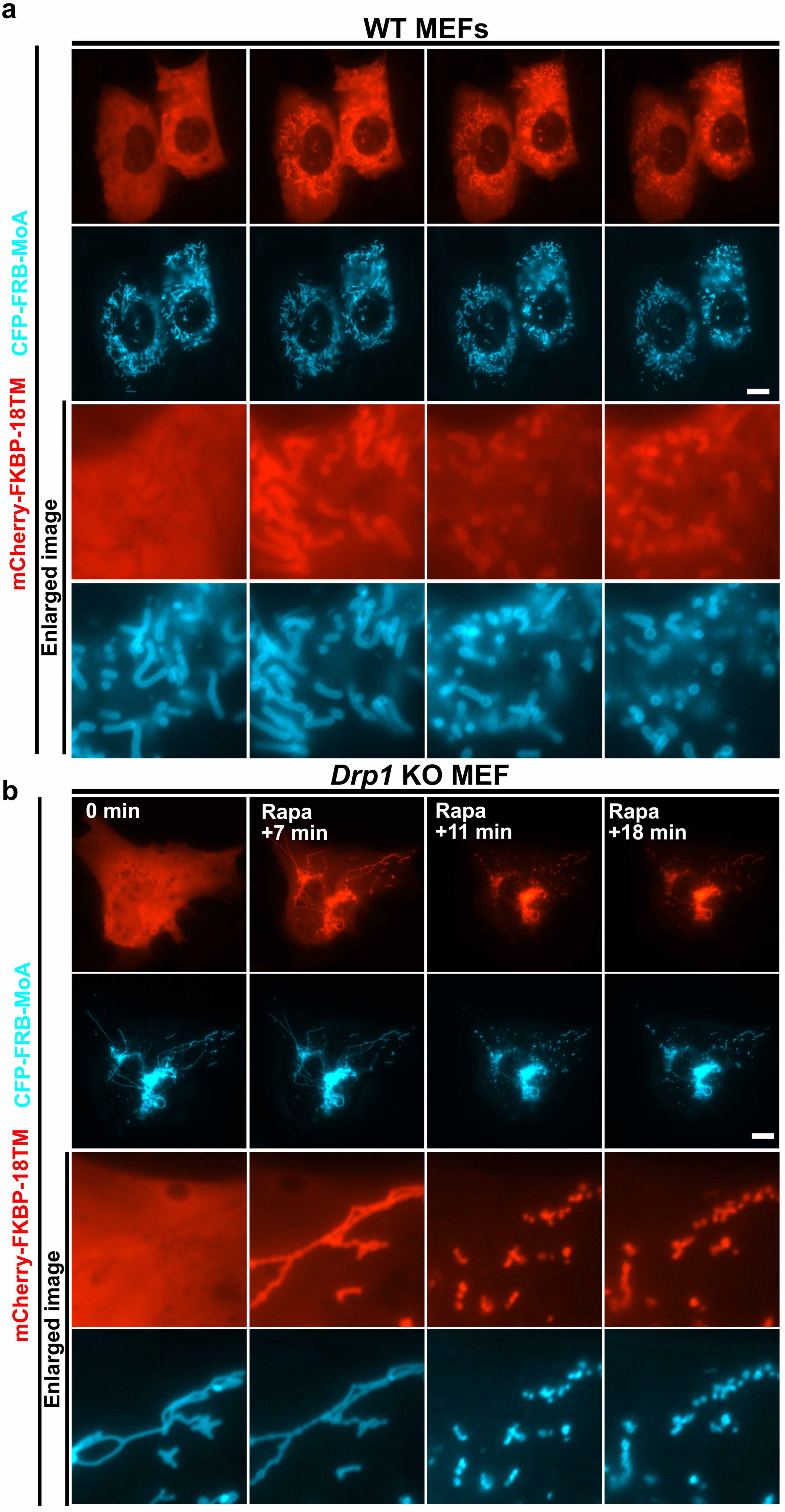

### Supplementary Figure 10

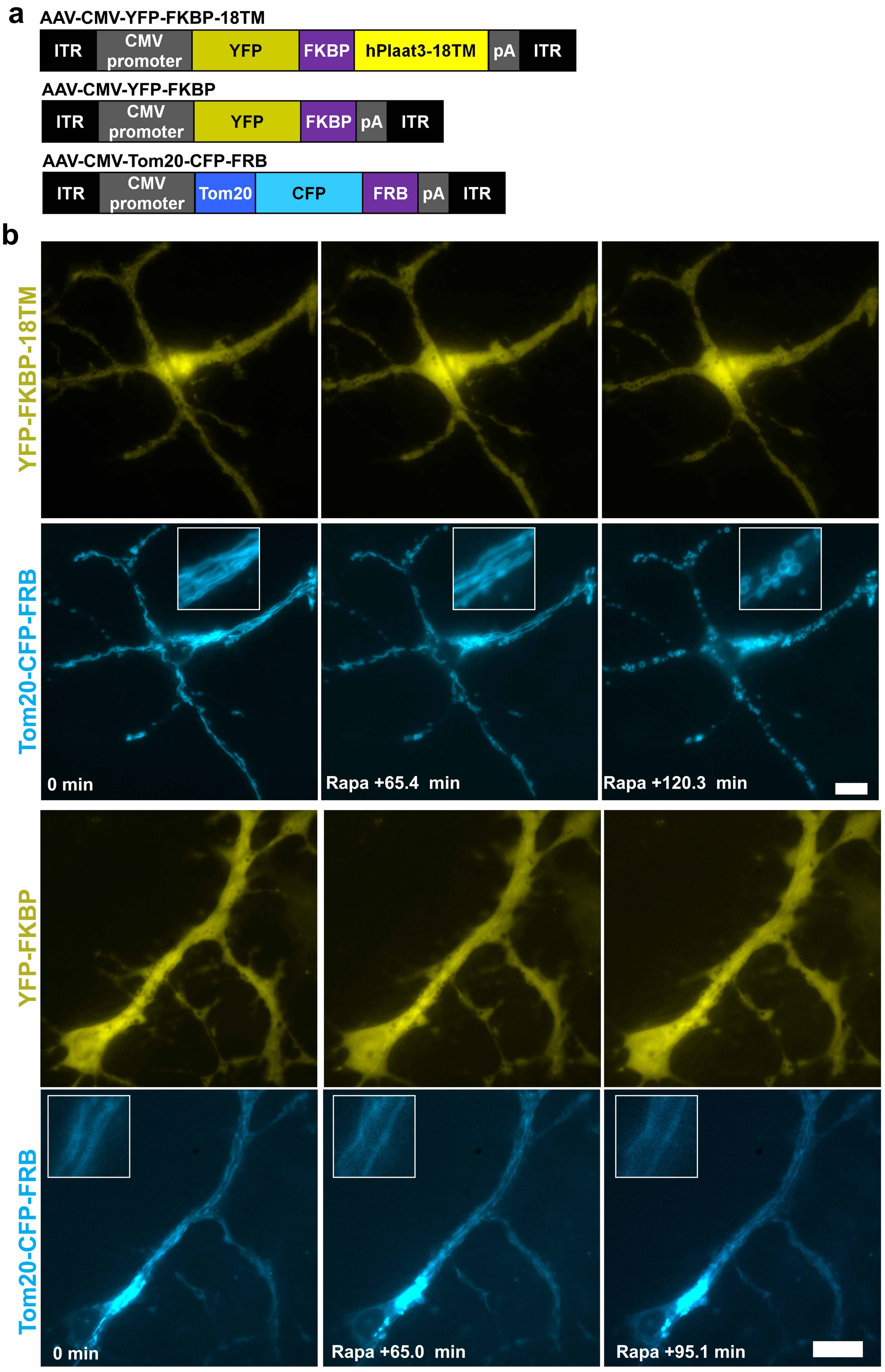
