## Supplementary Information for "Defunctionalizing Intracellular Organelles with Genetically-Encoded Molecular Tools Based on Engineered Phospholipase A/Acyltransferases (PLAATs)"

Contents:

Supplementary Experiments 1 and 2
Supplementary Reference
Figure Legends for Supplementary Figures 1-10
Supplementary Movies 1-8

**Supplementary Experiment 1**

**Spontaneous and stress-induced translocation of PLAAT3**

In the cell viability assay, we became aware that FL and FL-LD basically exist in soluble forms but also exhibited local accumulation in most of cells (**Fig. 1e**, insets). In contrast, expression of 18TM did not elicit these punctate signals, suggesting that CT domain was responsible for inducing this punctuation. Though it has been reported that membrane-damage reagents induced murine PLAAT3 translocation onto organellar membranes[1], our result suggested that PLAAT3 may be spontaneously relocalized to organellar membrane by cellular stress as well. Thus, we further examine the property of damage-dependent translocation of 18TM. At first, we used l-Leucyl-l-Leucine methyl ester (LLOME) which was used as a positive control to directly destabilize organellar membrane[1]. After COS-7 cells were transfected with plasmids expressing YFP, YFP-FL, or YFP-18TM (or their mCherry versions) and CFP-MoA, we observed translocating to mitochondria after treatment with 1mM of LLOME or other cellular stress agents. LLOME induced co-localization of mCherry-FL and CFP signals on mitochondria in 26.7% of transfected cells (**Supplementary Fig. 7a, b**). We also observed that mCherry and mCherry-18TM was not recruited to mitochondria in almost all transfected cells (**Supplementary Fig. 7a**, top and bottom panels, **b**). Then we tried addition of stress agents such as oxidative stress ($H_2O_2$) and hyperosmotic stress (high concentration of sucrose or NaCl) to the same culture condition. Two hours after $H_2O_2$ treatment (final concentration: 100 and 500 μM), YFP-FL showed co-localization with CFP-MoA, but co-localization with CFP was not detected in YFP-alone control cells at all (**Supplementary Fig. 7c, d**). In contrast, YFP-18TM was not co-localized with mitochondrial CFP signals. Results showing a similar trend were obtained in hyperosmotic stress condition (**Supplementary Fig. 7e-h**). Though sucrose addition at 500 mM induced mitochondrial translocation of 18TM only in 12.6% of transfected cells, the other hyperosmotic condition hardly elicited it (**Supplementary Fig. 7f, h**). In summary, FL was translocated to mitochondria upon oxidative stress or hyperosmolarity challenge, but 18TM was largely not. These stress-induced recruitments were observed in cells transfected with LD mutant of FL, but not in ones transfected with mCherry or mCherry-18TM-LD (**Supplementary Fig. 8a-d**). These results indicate that oxidative and hyperosmotic stress in addition to direct membrane damage can induce relocation of PLAAT3 onto mitochondria, which was independent of PLA activity (**Supplementary Fig. 7i**), while18TM does not show spontaneous or stress-induced relocalization.

**Supplementary Experiment 2**

**Step-wise development of 18TM-induced mitochondrial deformation**

To address fine morphological changes following 18TM recruitment, we analyzed serial fluorescent images. We used COS-7 cells co-transfected with mCherry-FKBP-18TM and CFP-FRB-MoA. At first, blebbing occurred in initially tubular mitochondria 2-7 minutes after rapamycin treatment (**Fig. 2a-c**, ROI-1, ROI: region of interest). Then, blebs were swollen and exhibited pearling-like deformation about 9 minutes after rapamycin treatment. Subsequently, swelling went on and each bleb was finally torn off, meaning that mitochondria was fully fragmented and took round shape (**Fig. 2a-c**, ROI-2). A part of round forms of mitochondria shrank and about 36 minutes after rapamycin addition, C-shaped mitochondria were seen (**Fig. 2a-c**, ROI-3).

### Supplementary Figures

**Supplementary Figure 1.**

**a** Sequence alignment of amino acids of human PLAAT family proteins. PLAAT1: NP_065119, PLAAT2: NP_060348, PLAAT3: NP_009000, PLAAT4: NP_004576, and PLAAT5: NP_ 473449. Pro: Proline-rich domain, LRAT: lecithin-retinol acyltransferase, TM: putative transmembrane domain, CT: c-terminal domain. LRAT and TM domains were assigned based on information of UniProtKB. Sequences were aligned by Clustal Omega. **b** Fluorescent images of cells expressing mCherry-FKBP-PLAAT1, PLAAT2, PLAAT4, PLAAT4-dTM and PLAAT5 along with CFP-FRB-MoA (mitochondria outer membrane anchor protein) before and after rapamycin treatment. mCherry-FKBP was used as a negative control. Insets indicate high magnification images. Scale bar = 10 μm. Rapa: rapamycin.

**Supplementary Figure 2. C-terminal fusion of a dimerizing unit to PLAAT3-FL abrogated mitochondrial deformation activity**

A full-length PLAAT3 was fused C-terminally with FKBP-YFP. A CID deformation assay was conducted in COS-7 cells, but mitochondria deformation was not induced 30 mins rapamycin. Areas marked with dashed white boxes were enlarged on the right side. Scale bar = 10 μm. Rapa: rapamycin.

**Supplementary Figure 3. Subcellular localization of PLAAT3 truncation mutants**

**a-c** Fluorescent microscopy of co-localization of FL, dCT, dTM, 2CT, and 18TM with PEX3-YFP (**a**), CFP-SEC61B (**b**) and CFP-MoA (**c**). Correlation of co-localization was analyzed by line scanning. Fluorescent intensities were analyzed along white arrows and normalized by average of intensities at all points. Cells were analyzed at day1 post-transfection. Subcellular localization of FL and its mutants was summarized in Fig. 1c. Scale bar = 10 μm.

**Supplementary Figure 4. Subcellular localization and mitochondrial deformation activity of PLAAT3-15TM, 19TM and 20TM**

**a-c** Representative images of fluorescence of COS-7 cells expressing 15TM (**a**), 19TM (**b**) and 20TM (**c**) before and after rapamycin treatment. These truncation mutants were localized in cytosol and did not show apparent mislocalization of any membrane organelle (**a-c**, upper panels). Activity of inducible deformation of mitochondria was seen in 19TM- and 20TM-expressing cells, but not in 15TM-expressing cells (**a-c**, lower panels). Areas marked with dashed white boxes were enlarged on the right side. Scale bar = 10 μm. Rapa: rapamycin.

**Supplementary Figure 5. Localization of PLAAT3-FL and -18TM on other membrane-bound organelles**

**a-e** Subcellular localization of FL and 18TM on Golgi apparatus (Giantin-CFP) (**a**), lysosomes (LAMP1-CFP) (**b**), autophagosomes (mCherry-LC3) (**c**), endosomes (mCherry-Rab5) (**d**), and nucleus (mCherry-Lamin A) (**e**). To detect autophagosomes, cells were treated with chloroquine (final concentration: 100 μM) for one overnight. YFP-alone sample reflects cytosolic localization pattern. No correlation between each organelle marker signal and signal from FL or 18TM. Scale bar = 10 μm.

**Supplementary Figure 6. PLAAT3-18TM did not reduce peroxisomal number**

Effect of FL and 18TM expression on peroxisomal number. Cells were transfected with plasmids expressing YFP (negative control), YFP-FL, and YFP-18TM and imaged at 48 hrs after transfection. FL expression reduced dot signals from mSca-peroxi, but 18TM expression did not. Scale bar = 10 μm.

**Supplementary Figure 7. Relocation of PLAAT3-FL and -18TM upon oxidative or hyperosmotic stress**

**a** Observation of relocation of FL and 18TM 1 hr after treatment with membrane damage agent (1mM LLOME). **b** Quantification of **a**. n = 405, 305 and 361 cells from left to right; analyzed from three individual experiments. Fluorescent images of cells treated with oxidative stress (100 μM or 500 μM of $H_2O_2$) or hyperosmotic stress (200 mM or 500 mM of Sucrose, 50 mM or 100 mM of NaCl) for 2 hrs. **d**, **f**, **h** Quantification of **c**, **e** and **g**. n = 222, 240, 213, 255, 192 and 252 cells from left to right; analyzed from three individual experiments (**d**). n = 373, 379, 273, 265, 277 and 216 cells from left to right; analyzed from three individual experiments (**f**). n = 297, 264, 206, 273, 229 and 234 cells from left to right; analyzed from three individual experiments (**h**). Percentage of cells with recruitment of mCherry, mCherry-FL, and mCherry-18TM on mitochondria was calculated.

**i** Summary of spontaneous recruit and stress-induced recruit of FL, FL-LD, 18TM and 18TM-LD. Data from Fig. 1e and data from Supplementary Fig. 6 were integrated to **i** for summary. Insets indicate high magnification images. Error bars indicate means ± s.d.. Scale bar = 10 μm.

**Supplementary Figure 8. Stress-induced relocation was independent on PLA activity**

**a-d** Fluorescent analysis of recruitment of FL-LD and 18TM-LD in 1mM-LLOME-treated cells (**a**), 500μM-$H_2O_2$-treated cells (**b**), 500mM-sucrose-treated cells (**c**), and 100mM-NaCl-treatd cells (**d**). COS-7 cells were transfected with plasmids expressing mCherry, mCherry-FL-LD, and mCherry-18TM-LD. mCherry-alone sample is negative control. Scale bar = 10 μm.

**Supplementary Figure 9. PLAAT3-18TM-mediated fragmentation was independent of mitochondrial fission exerted by DRP1**

**a**, **b** Representative images of WT (**a**) or *Drp1* KO (**b**) MEFs expressing mCherry-FKBP-18TM and CFP-FRB-MoA before and after rapamycin treatment. In *Drp1* KO MEFs, only elongated forms of mitochondria were seen. In both cell types, mitochondrial deformation by 18TM was induced. Scale bar = 10 μm. Rapa: rapamycin.

**Supplemental Figure 10. Induced mitochondrial deformation following AAV-mediated delivery of 18TM**

**a** Schematic diagram of AAV transgenes. Three types of AAVs were generated and infected to mouse primary hippocampal neurons. The fist type of AAV expresses YFP-FKBP-18TM, second expresses YFP-FKBP-alone and third expresses Tom20-CFP-FRB (mitochondrial outer membrane anchor). Expression of all genes was driven by CMV promoter. AAV-CMV-YFP-FKBP was used as a negative control. ITR: inverted terminal repeat. Viruses were added to neurons at 6 DIV at MOI = 40,000. Two days after infection, cells were analyzed. **b** Representative images of mitochondrial morphology in neurons. Mitochondrial deformation after 18TM translocation was induced in YFP-FKBP-18TM-expressing neurons. Scale bar = 10 μm. Rapa: rapamycin.

**Supplementary Movies**

**Supplementary Movies 1.** Inducible mitochondrial deformation using PLAAT3-FL. COS-7 cells were transfected with mCherry-FKBP-PLAAT3-FL and CFP-FRB-MoA. The upper panels of **Fig. 1** were extracted as representative images based on this movie. Rapamycin was added at t = 5 mins. Scale bar = 10 μm.

**Supplementary Movies 2.** A lipase dead mutant of PLAAT3 did not induce mitochondrial deformation. COS-7 cells were transfected with mCherry-FKBP-PLAAT3-FL-LD and CFP-FRB-MoA. The lower panels of **Fig. 1** were extracted as representative images based on this movie. Rapamycin was added at t = 5 mins. Scale bar = 10 μm.

**Supplementary Movies 3.** More detailed morphological changes of mitochondria upon a CID-18TM recruitment. COS-7 cells were transfected with mCherry-FKBP-18TM and CFP-FRB-MoA. The images of **Fig. 2a** were extracted as representative images based on this movie. Rapamycin was added at t = 1 min. Scale bar = 10 μm.

**Supplementary Movies 4.** Loss of membrane potential following the 18TM-medtiated deformation. COS-7 cells were transfected with YFP-FKBP-18TM and CFP-FRB-MoA. The images of **Fig. 3a** were extracted as representative images based on this movie. Rapamycin was added at t = 9 mins. Scale bar = 10 μm.

**Supplementary Movies 5.** Su9-CFP leakage following the 18TM-mediated deformation. COS-7 cells were transfected with YFP-FKBP-18TM, Su9-CFP, and mCherry-FRB-MoA. The images of **Fig. 4a** were extracted as representative images based on this movie. This movie shows only CFP (blue) and mCherry (red) channels. Scale bar = 10 μm.

**Supplementary Movies 6.** Su9-CFP did not leak out of mitochondria in 18TM-LD-expressing cells. COS-7 cells were transfected with YFP-FKBP-18TM-LD, Su9-CFP, and mCherry-FRB-MoA. The images of **Fig. 6c** were extracted as representative images based on this movie. This movie shows only CFP (blue) and mCherry (red) channels. Scale bar = 10 μm.

**Supplementary Movies 7.** Dissipation of matrix-resident proteins of peroxisomes following the 18TM recruitment. HeLa cells were transfected with YFP-FKBP-18TM, PEX3-CFP-FRB, and mSca-Peroxi. The images of **Fig. 6a** were extracted as representative images based on this movie. Rapamycin was added at t = 1 min. Scale bar = 10 μm.

**Supplementary Movies 8.** mSca-Peroxi proteins were not dissipated following the 18TM-LD recruitment. HeLa cells were transfected with YFP-FKBP-18TM-LD, PEX3-CFP-FRB, and mSca-peroxi. The images of **Fig. 6b** were extracted as representative images based on this movie. Rapamycin was added at t = 1 min. Scale bar = 10 μm.
